# Integrated technology platform for accelerated discovery of antiviral antibody therapeutics

**DOI:** 10.1101/2020.05.12.090944

**Authors:** Pavlo Gilchuk, Robin G. Bombardi, Jesse H. Erasmus, Qing Tan, Rachel Nargi, Cinque Soto, Peter Abbink, Todd J Suscovich, Lorellin A. Durnell, Amit Khandhar, Jacob Archer, Jenny Liang, Mallorie E. Fouch, Edgar Davidson, Benjamin J. Doranz, Taylor Jones, Elise Larson, Stacey Ertel, Brian Granger, Jasmine Fuerte-Stone, Vicky Roy, Thomas Broge, Thomas C. Linnekin, Caitlyn H. Linde, Matthew J. Gorman, Joseph Nkolola, Galit Alter, Steven G. Reed, Dan H. Barouch, Michael S. Diamond, James E. Crowe, Neal Van Hoeven, Larissa Thackray, Robert Carnahan

## Abstract

The emergence and reemergence of highly virulent viral pathogens with pandemic potential has created an urgent need for accelerated discovery of antiviral therapeutics. Antiviral human monoclonal (mAbs) are promising drug candidates to prevent or treat severe viral diseases, but the long timelines needed for discovery limits their rapid deployment and use. Here, we report the development of an integrated sequence of technologies incorporating advances in single-cell mRNA sequence analysis, bioinformatics, synthetic biology, and high-throughput functional analysis that allowed us to discover highly potent antiviral human mAbs and validate their activity *in vivo* at an unprecedented scale, speed, and efficiency. In a 78-day study, modeling deployment of a rapid response platform to an outbreak, we isolated >100 individual Zika virus (ZIKV) specific human mAbs, assessed their function, identified 29 broadly-neutralizing mAbs, and verified therapeutic potency of lead candidates with antibody-encoding mRNA formulation and/or IgG protein delivery in mice and nonhuman primates. Our work provides a roadmap for the rapid antibody discovery programs against viral pathogens of global concern.

## Main

Human monoclonal antibodies (mAbs) are increasingly being considered as therapeutic countermeasures for viral infectious diseases. In recent years, advances in human B cell isolation and antibody variable gene sequencing techniques have led to the identification of large numbers of therapeutic mAb candidates against many life-threatening viral pathogens. These targets include antigenically variable viruses such as human immunodeficiency virus^1^ and influenza virus^2^, newly emerging pathogens with high epidemic potential including Ebola virus^3^, Marburg virus^4^, Zika virus (ZIKV)^5-7^, Lassa virus^8^, Middle East respiratory syndrome coronavirus (MERS-CoV)^9^, poxviruses^10^, Nipah virus^11^ and many other medically important viruses. Over 25 antiviral human mAbs are now being evaluated as therapeutics in clinical trials^12,13^.

Large human epidemics of zoonotic diseases are occurring on a regular basis. For example, Ebola virus reportedly caused 28,646 cases of disease and 11,323 deaths in the 2013–2016 epidemic in West Africa^14^, the 2015-2016 ZIKV epidemic resulted in millions of infections in new geographic areas^15,16^ and recently, a novel, highly transmissible coronavirus, 2019-nCoV emerged in China and spread globally such that by February 27^th,^ 2020, 82,294 cases and 2,747 deaths have been reported (WHO: https://www.who.int/emergencies/diseases/novel-coronavirus-2019/situation-reports). These events highlight the potential of viral infections to cause global health emergencies and the need for rapid response platforms to accelerate development of medical countermeasures. Several obstacles impede progress in widespread application of antiviral mAb therapies in outbreak scenarios. A principal impediment to deployment of human antiviral mAbs is the difficulty in predicting which pathogen will cause an epidemic in the short term, and the long timeline needed for human mAb discovery and verification of therapeutic potency.

The goal of this study was to develop and demonstrate the capability of an integrated platform for accelerated discovery of potent human antiviral mAbs that are suitable for therapeutic development. As a model, we used the reemerging pathogen ZIKV that is now endemic in multiple continents^17^ and potentially could be controlled with a neutralizing mAb treatment^5,16,18,19^. We designed an integrated workflow that could accomplish both discovery and validation of protective efficacy in small animals and nonhuman primates (NHP) in less than 90 days. The objectives were to execute rapidly a sequence of tasks including virus stock production, target-specific antibody genes rescue, bioinformatics analysis, mAb gene synthesis, mAb functional *in vitro* analysis, validation of mAb *in vivo* activity using IgG protein and RNA-encoded mAb delivery in mouse model, and proof-of-principle NHP protection studies, as outlined in **Fig. 1**. The workflow implemented high-throughput and redundant approaches, with the goal of streamlining mAb discovery and building in orthogonal approaches to overcome or mitigate failure of critical experimental processes.

**Fig. 1.**
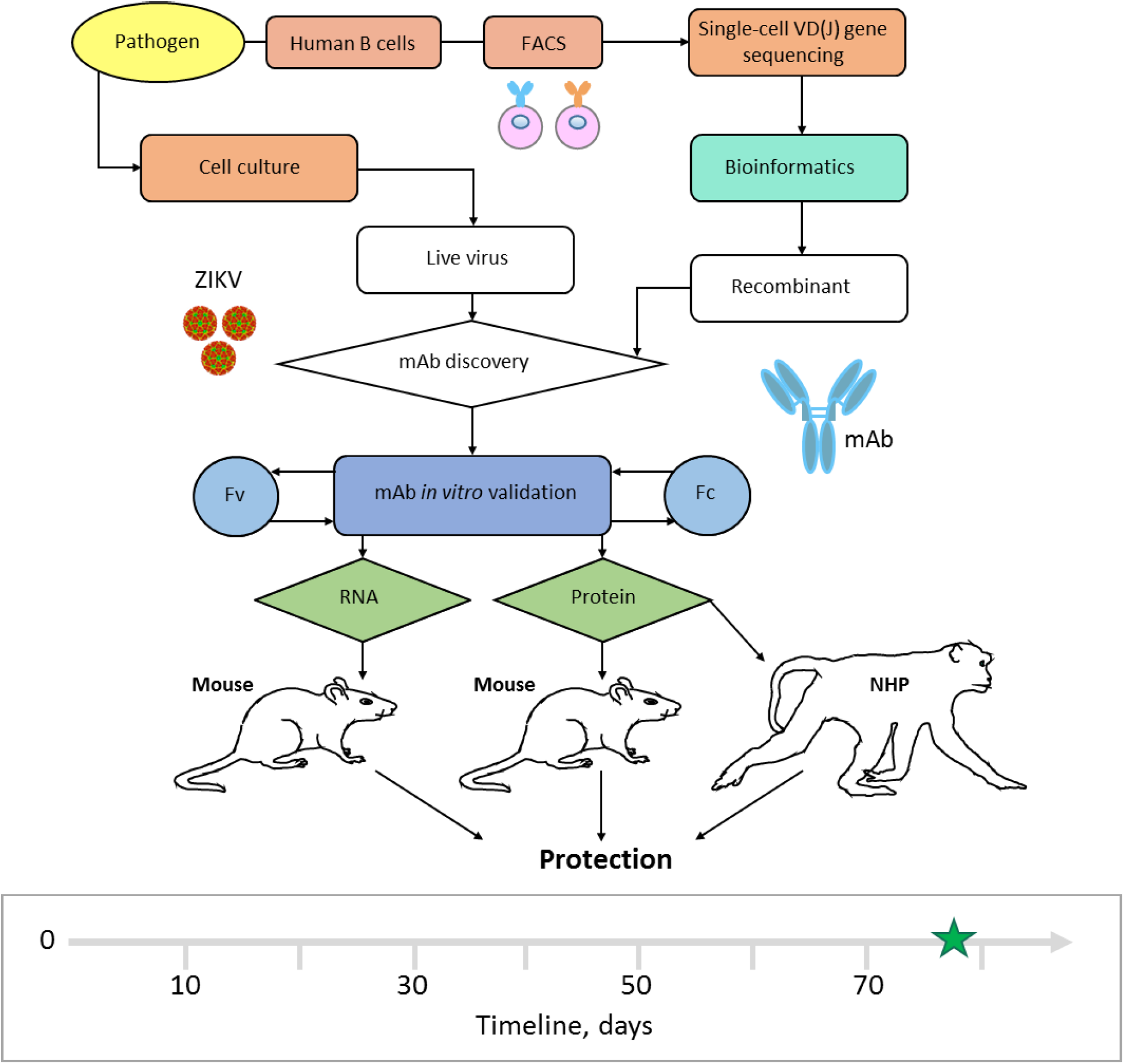
Integrated technology workflow for rapid discovery of anti-viral human mAbs. A cartoon of proposed integrated technology workflow that incorporated pathogen production, target-specific B cell isolation, single-cell VD(J) genes sequence analysis, bioinformatics analysis, mAb production, mAb *in vitro* validation, mAb-encoding RNA synthesis, and mAb *in vivo* validation using protein and nucleic acid delivery technologies. Timeline to identify and validate protective mAbs against ZIKV is indicated in the timeline chart, where day 0 is designated as a start point, and day 90 was the projected endpoint for this study. Green star (day 78) depicts the completion time.

## Results

### Virus stock production

We used a contemporary ZIKV strain of the Asian lineage (Brazil, Paraiba 2015) and a historical ZIKV strain of the African lineage (Dakar, Senegal 1984) to account for genetic diversity and ultimately breadth of protection. To model ZIKV infection in mice, we also used a mouse-adapted isolate of ZIKV Dakar (ZIKV Dakar MA), which has a single gain-of-function mutation in the viral NS4B protein that overcomes mouse innate immune restriction^20^. To model production of a high-titer virus stock for an unknown pathogen, we evaluated the growth of ZIKV in a panel of cell lines that are commonly used for virus propagation. High levels of virus replication were observed in Vero, U20S, A549, and Huh7 cell lines, as determined by quantitative reverse-transcription PCR (qRT-PCR) (**Supplementary Fig. 1a**), showing the utility of this approach for rapid identification of a permissive cell line(s). Identifying suitable conditions for large-scale production of the virus for subsequent *in vitro* and *in vivo* studies (**Supplementary Fig. 1b-c**) was accomplished in 21 days. Of note, the virus production was performed simultaneously and in a parallel with initial mAb discovery steps, and therefore, the virus was obtained when it was needed for the neutralizing activity screening step with purified mAbs, as detailed below.

### Antibody variable gene sequence analysis and bioinformatic processing

To model a rapid response to a previously known pathogen, we used a mAb discovery approach that assumed the envelope protein (ZIKV E protein) on the virion surface is a key protective antigen^5,6,21,22^. We used memory B cells from previously infected human subjects as the source for antibody discovery (**Supplementary Table 1**). The frequency of ZIKV E-specific memory B cells in most human subjects with prior ZIKV infection is low^23^. Therefore, we pooled peripheral blood mononuclear cells (PBMCs) from a cohort of several subjects with previous exposure to ZIKV that were pre-screened for E-specific memory B cell responses (**Supplementary Fig. 2**). B cells were enriched using antibody-coated magnetic beads and stained with phenotyping antibodies and soluble recombinant ZIKV E protein (see *Methods*). Antigen-labeled IgG class-switched memory B cell-E complexes were isolated in bulk using fluorescence activated cell sorting (FACS) (**Fig. 2a**). To optimize the breadth of our mAb panel for diverse epitope specificities, we used several B cell isolation approaches that were based on direct or indirect labeling with ZIKV E protein, which resulted in generation of three independent sub-panels of mAbs (**Supplementary Table 2**). Overall, > 5,000 E-labeled B cells were sorted.

**Fig. 2.**
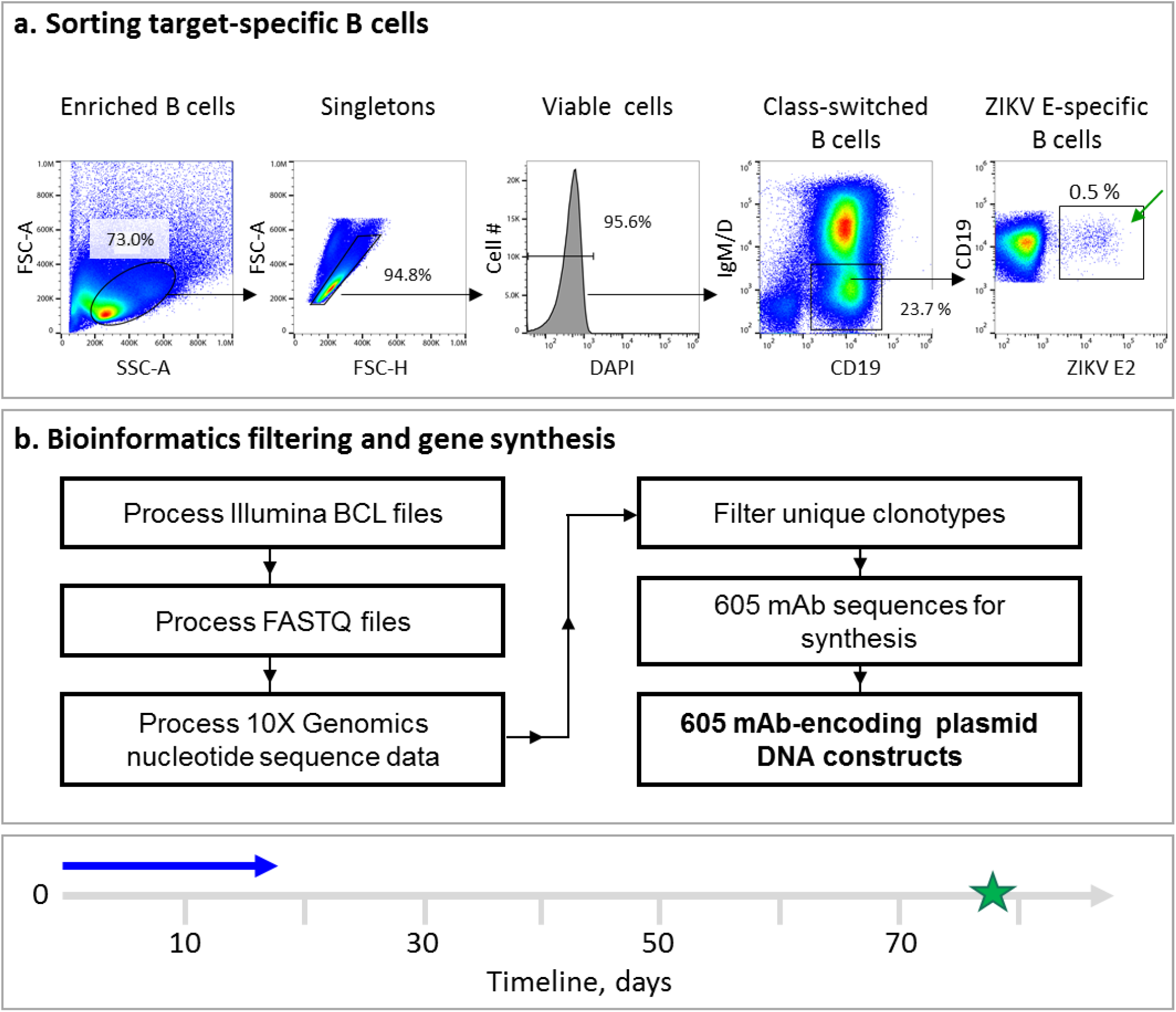
Rescue and sequence analysis of antibody genes from ZIKV E antigen-specific human B cells. **a**, Flow cytometric identification of target-specific memory B cells after labeling of magnetically enriched total B cells with ZIKV E antigen. Target-specific B cells were labeled with biotinylated recombinant E protein and detected by fluorochrome-conjugated streptavidin as indicated. Green arrow indicates FACS-sorted cells. Representative data of four experiments is shown. **b**, Bioinformatics filtering steps to select mAb sequences for synthesis. Timeline to identify sequences of target mAbs and obtain cDNA constructs for recombinant mAb expression is indicated with blue arrow in the timeline chart.

To rescue the paired heavy and light chain antibody variable gene sequences from sorted B cells, we used a commercial single-cell encapsulation automated system (Chromium Technology; 10x Genomics Inc). A feature of this sequencing approach is the unprecedented scale of paired antibody variable gene analysis, which is achieved by encapsulation of single B cells via droplet microfluidics^24^. Approximately 800 sorted antigen-labeled B cells from the single specimen of pooled human cells were subjected to library preparation and sequencing using the Chromium Controller device immediately after cell isolation by FACS. The remaining sorted cells were bulk-expanded in culture, and then the antibody variable genes for single cells were sequenced using Chromium Technology.

Using bioinformatics sieving, we down-selected the total pool of possible sequences to choose 598 paired antibody heavy and light chain variable gene sequences for further evaluation (**Fig. 2b**). The filtered panel included mAbs of defined IgG isotypes, selecting the most somatically mutated individual clone in each antibody clonotype (*i.e.*, the clone with the most mutations compared to the inferred germline gene segments in the lineage of related sequences). Clonotypes were identified based on the presence of identical inferred germline V and J gene assignments and identical amino acid sequences of the CDR (See *Methods*).

Synthetic cDNA constructs based on the sequences of all 598 selected ZIKV mAb candidates were synthesized using a rapid high-throughput cDNA synthesis platform (Twist Bioscience) and cloned into an IgG1 monocistronic expression vector for mammalian cell culture mAb secretion. This vector contains an enhanced 2A sequence and GSG linker that allows simultaneous expression of mAb heavy and light chain genes from a single construct upon transfection^25^.

In summary, starting from human subject samples, in 17 days we produced recombinant IgG expression vectors for nearly 600 human mAbs from single cells representing independent genetic clonotypes that were ready for expression and functional validation (**Supplementary Table 2**). The data demonstrated capability for rapid target-specific B cell selection and large-scale human antibody discovery, for a setting in which a known protective viral antigen is available.

### High-throughput mAb production and functional screening

We adopted high-throughput assays for rapid production, purification, and functional analysis of mAbs from small sample volumes (designated here as “micro-scale”) to identify lead mAb candidates for *in vivo* studies. To produce mAbs, we performed transfection of Chinese hamster ovary (CHO) cell cultures using ∼1 mL cell culture volume per antibody, and mAbs subsequently were purified using a micro-scale parallel purification approach. Of the 598 mAb cDNAs tested, 475 were produced successfully as recombinant IgG proteins, with a median yield of ∼29 µg of purified antibody from a single 1 mL culture (**Fig. 3a; Supplementary Table 2**).

**Fig. 3.**
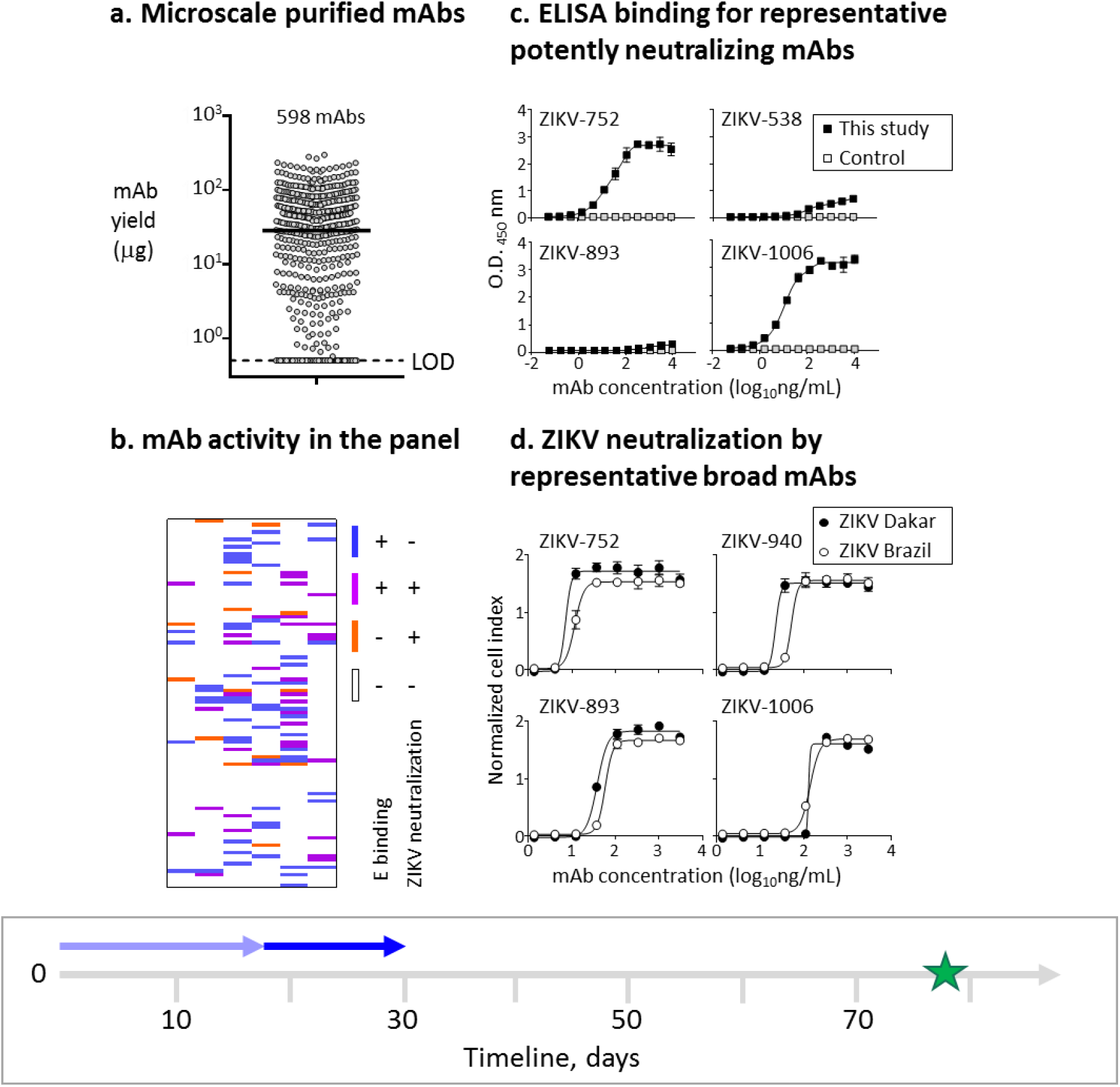
Rapid mAb production and screening to identify lead candidates for *in vivo* testing. **a**, Micro-scale expressed and purified mAbs. Dots indicate average concentration of individual mAbs from assay duplicates, and median mAb yield is shown with horizontal line. LOD – limit of the detection. **b**, Relationships between binding and neutralizing activities of individual mAbs of the panel shown as a heatmap. Binding to E protein was determined by ELISA and neutralizing activity was measured using RTCA from mAbs purified as in (a). **c**, ZIKV E protein binding by potently neutralizing mAbs representing three distinct epitope binding groups. MAb rRSV-90, which is specific to the unrelated respiratory syncytial virus (RSV) fusion protein (F) antigen, served as a control. **d**, Neutralization of ZIKV Brazil or Dakar strain viruses by representative cross-neutralizing mAbs from (c), as determined using RTCA. Data shown indicate the mean ± SD of assay triplicates in (c, d), and represent at least two independent experiments. Timeline to identify mAb candidates for *in vivo* testing is indicated with a blue arrow in the timeline chart.

Antibodies can mediate protection by diverse mechanisms, including direct virus neutralization and/or engagement of innate immune cells via their Fc receptors (FcRs)^26,27^. Potent neutralization and Fc effector functions are desirable mAb features that can be used to prioritize candidate mAbs for *in vivo* testing. For rapid identification of neutralizing mAbs within the large panel we expressed, we used a real-time cell analysis (RTCA) cellular impedance assay. RTCA assesses kinetic changes in cell physiology, including virus-induced cytopathic effect (CPE), which offers a generic screening approach for mAb neutralizing activity against cytopathic viruses^28,29^. Our comparative side-by-side study showed that the RTCA performed similarly to a conventional focus-reduction neutralization test (FRNT) to identify neutralizing mAbs against ZIKV (**Supplementary Table 2**). However, RTCA offered an advantage for the large mAb panel analysis due to its speed and automation — 24 to 36 hours for the initial qualitative screening of neutralizing activity, and ∼48 hours for determination of half maximal inhibitory concentration (IC_50_) values and ranking of ZIKV mAbs by neutralizing potency (**Supplementary Fig. 3**). Approximately 15% percent (92 of 598) of the mAbs bound strongly to recombinant ZIKV E protein as measured by ELISA, and ∼8% (48 of 598) of the mAbs neutralized ZIKV (**Fig. 3b-d; Supplementary Table 2**). Fourteen additional ZIKV-specific mAbs were identified only by neutralization, as they did not show detectable binding to the recombinant E antigen in the solid-phase by ELISA, even though their B cells were sorted with the same antigen in solution. Thus, using a combination of binding and neutralization assays, we validated that at least 106 of the mAbs from the panel were ZIKV-specific (**Supplementary Table 2**).

The subjects we studied likely had been infected with a strain of Asian lineage of ZIKV, since they contracted the disease during the recent outbreak in South America (**Supplementary Table 1**). Twenty-nine mAbs of the panel possessed neutralizing activity against viruses that were antigenically homologous (Brazil strain; Asian lineage) or heterologous (Dakar strain; African lineage) to the infecting strain (**Supplementary Table 2**). Twenty mAbs fully neutralized at least one of two tested viruses, with estimated IC_50_ values from RTCA data ranging from about 7 to 1,100 ng/mL, and 15 mAbs fully neutralized both viruses (**Fig. 4**). The three most potent mAbs by neutralization IC_50_ value (designated mAbs ZIKV-893, −752, and −940) neutralized both Brazil and Dakar ZIKV strains with IC_50_ values below 100 ng/mL but exhibited differential binding to recombinant E protein. ZIKV-752 bound E strongly, ZIKV-940 bound weakly, and binding of ZIKV-893 was not detected at the highest mAb concentration tested (10 µg/mL) (**Fig. 3c-d; 4**). Competition-binding analysis with the previously characterized ZIKV E-specific human mAbs ZIKV-117 (recognizes the E protein dimer-dimer interface in domain II on viral particles)^30^, ZIKV-116 (binds to the E protein domain III lateral ridge epitope)^31^, and ZIKV-88 (specific for the E protein fusion loop)^5^, showed that the three newly identified neutralizing mAbs ZIKV-893, - 752, and −940 recognized non-overlapping epitopes (**Fig. 4; Supplementary Table 2**). This conclusion was supported by epitope mapping data from loss-of-binding analysis to a ZIKV prM-E protein alanine scanning mutagenesis library expressed in cells^5^. Notably, these mAbs were identified from three different E-antigen sorting strategies (**Supplementary Table 2**). This finding highlighted the importance of study design in the antibody discovery workflow, which relied on concurrent independent strategies for B cell isolation conducted in parallel to identify potently neutralizing mAbs of diverse epitope specificity.

**Fig. 4.**
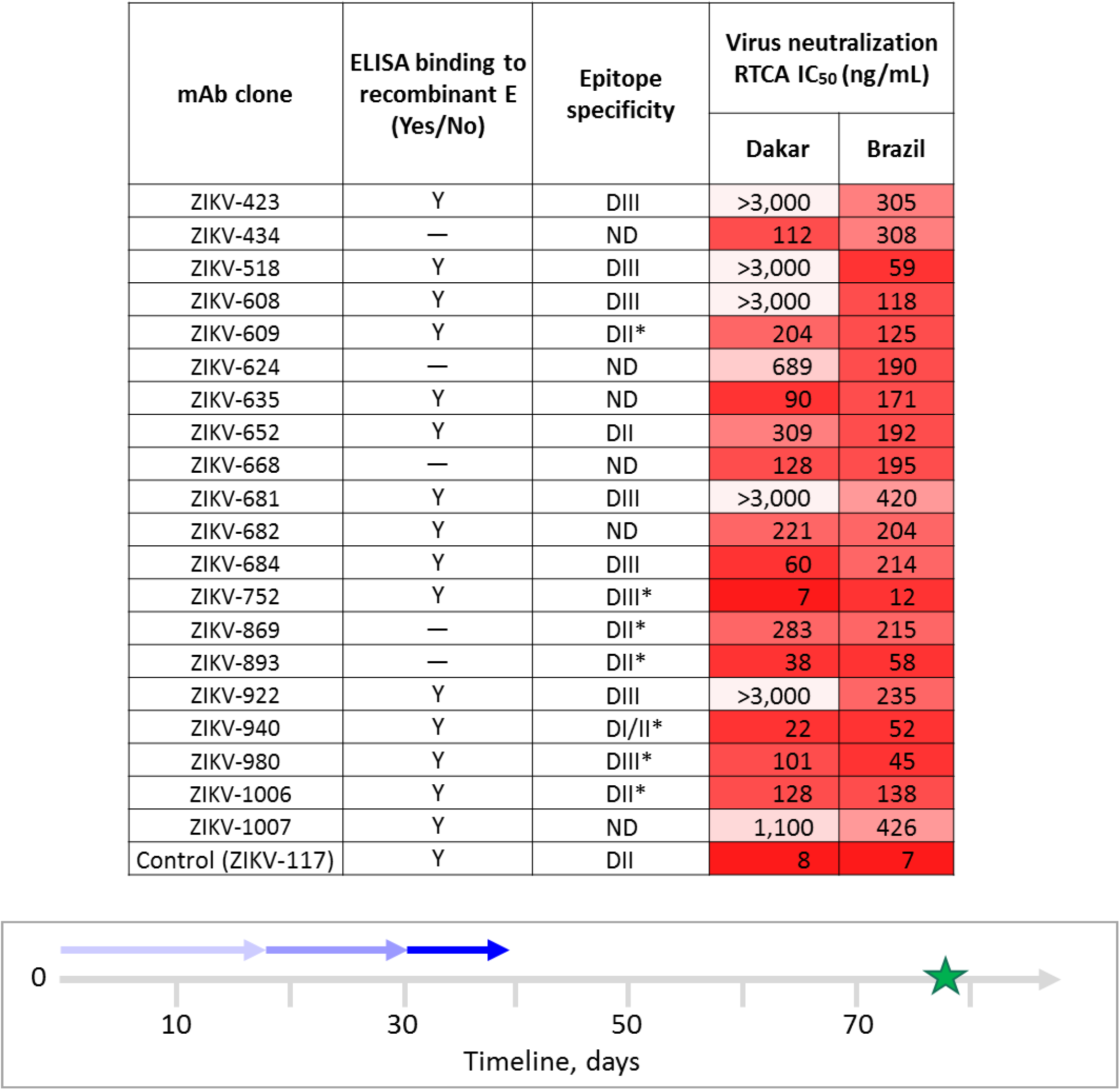
Binding and functional activities of lead human ZIKV mAb therapeutic candidates selected for in vivo testing. Twenty mAbs were selected as lead therapeutic candidates on the basis of neutralizing potency and breadth. The measured activities of mAbs are indicated with values and presented as a heatmap. The “*” symbol indicates epitope specificity that was mapped precisely by a shotgun mutagenesis assay assessing loss of binding of point mutant E proteins. Timeline to scale-up mAb protein and mAb-encoding RNA production for in vivo testing in mice is indicated with a blue arrow in the timeline chart.

To demonstrate that Fc-mediated function profile could be determined rapidly for this large mAb panel, we performed antibody-dependent cellular phagocytosis (ADCP), antibody-dependent neutrophil phagocytosis (ADNP), and antibody-dependent complement deposition assays in a 384-well format. These assays used immobilized ZIKV E protein and high-throughput flow cytometric analysis to determine the capacity of bound mAb to activate human effector cells *in vitro*. Many mAbs of the panel revealed a high capacity to engage FcRs (**Supplementary Table 2**), showing the utility of the approach for rapid identification of mAbs leveraging Fc function.

In summary, these data demonstrated the high performance of a suite of micro-scale assays in an integrated workflow and accomplished accelerated production and functional analysis of large panels of recombinant human antibodies to identify lead candidates for *in vivo* studies – all of which was accomplished in 13 days.

### IgG protein and RNA production for the lead therapeutic mAb candidates for *in vivo* studies

We selected 20 lead mAb candidates for *in vivo* testing. We administered the mAbs using two different delivery systems: 1) recombinant IgG protein expressed in mammalian cell culture, and 2) mRNA delivery of antibody genes to allow *in vivo* expression of antibodies from transient gene transfer. We initiated production of 20 recombinant IgG proteins in cultures scaled to obtain 3-7 milligrams of purified IgG. In parallel, the same 20 antibody variable gene sequences were cloned into a reading frame for human IgG1 and sub-cloned into plasmids containing an alphavirus replicon^32^ for *in vivo* study with nucleic acid delivery. To deliver the replicon, we chose a cationic nanostructured lipid carrier (NLC) formulation, in which RNA is bound to the nanoparticle surface by electrostatic association, allowing for rapid formulation^32^. Linearized DNAs were used as template to transcribe and post-transcriptionally cap replicon RNA at a 500 μg scale. The IgG protein preparation and RNA production and formulation both were accomplished in parallel in eight days.

### Protective efficacy of identified neutralizing mAbs in a lethal ZIKV challenge mouse model

We next used an established lethal, immunocompromised challenge model in mice for ZIKV^5,21^ to evaluate the protective capacity of the 20 lead candidate mAbs (**Fig. 4; Supplementary Table 3-4**) and identify lead mAbs for NHP studies. In prophylaxis experiments, C57BL6/J (B6) mice were treated via intraperitoneal (i.p.) injection with anti-IFN-alpha/beta receptor (Ifnar1) antibody and individual ZIKV mAbs on day −1. Candidate mAbs were compared to a potently protective human mAb ZIKV-117 that we described previously^5^. An irrelevant IgG1 isotype mAb (FLU-5J8, which is specific to the influenza A virus hemagglutinin protein^33^ or PBS served as a negative control. On day 0, mice were challenged via subcutaneous (s.c.) route with 10^3^ focus-forming units (FFU) of the ZIKV Dakar MA and monitored for 21 days for survival. High-dose prophylaxis (about 70 µg of mAb per mouse; ∼5 mg/kg mAb dose) provided complete protection from mortality for three of the five tested mAbs. MAb ZIKV-668 offered partial protection, whereas mAb ZIKV-434 and FLU-5J8 or PBS treatment did not protect (**Supplementary Table 4)**. Since multiple individual mAbs conferred full protection with high-dose mAb prophylaxis, we could not discriminate which mAb was the most effective *in vivo*. Therefore, we next tested a larger panel of ten neutralizing mAbs with a low-dose prophylaxis approach (using about 9 µg of mAb; 0.65 mg/kg mAb dose). Even at low doses, six of the ten human mAbs were detected readily in mouse blood as assessed on 2 dpi (**Fig. 5a**). As an early time-point correlate of mAb-mediated protection, we measured viremia from individual mice of each treatment group by qRT-PCR. All groups treated with ZIKV mAbs showed markedly (500-1,000 fold) reduced levels of virus measured as genome equivalents (GEQ) per 1 mL of serum by 2 dpi when compared to the control mAb FLU-5J8-treated group, which developed high viremia in all animals (**Fig. 5b**). This finding demonstrated efficient inhibition of viremia by a low dose of anti-ZIKV mAb in a prophylaxis setting.

**Fig. 5.**
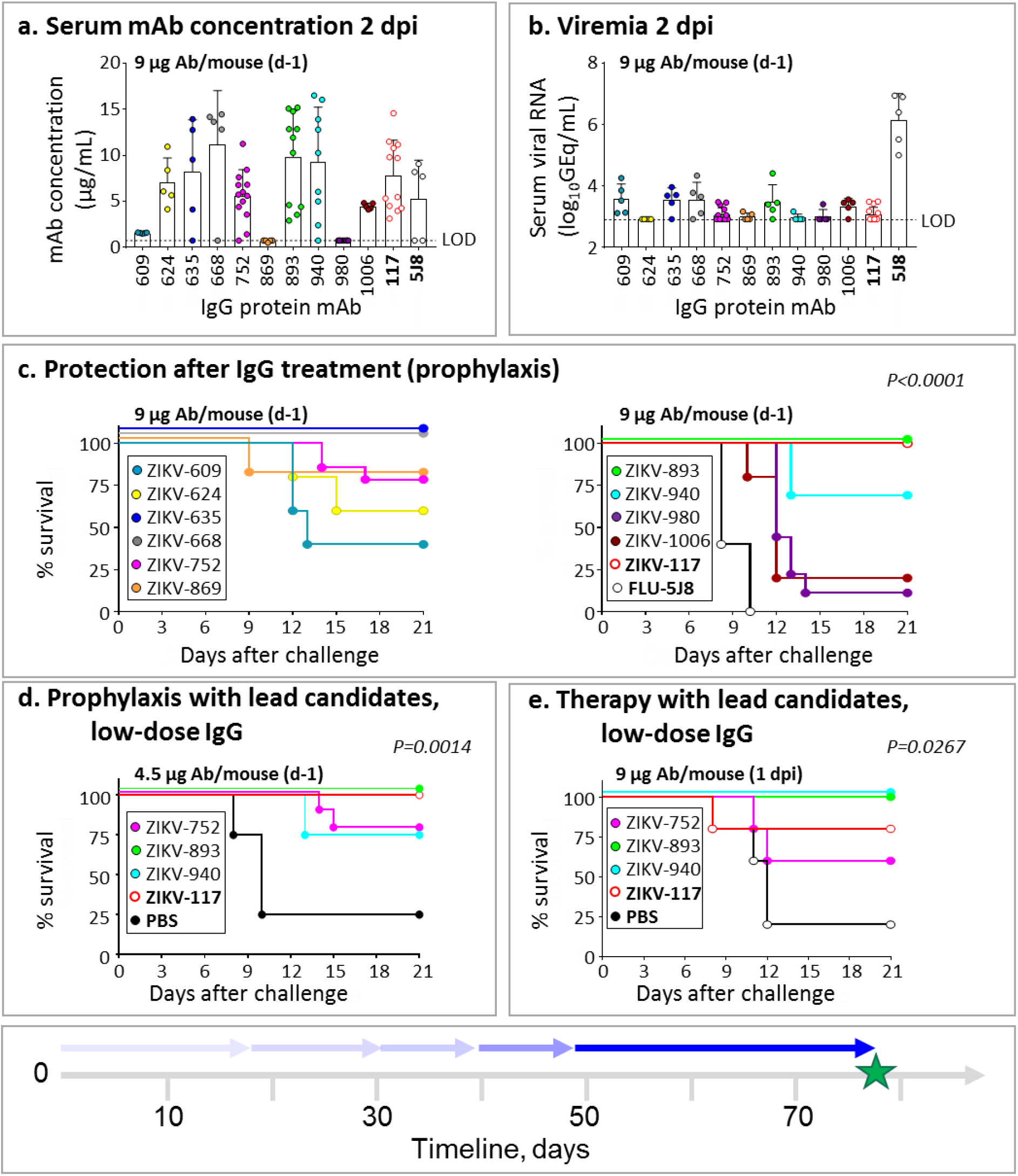
MAb protection against lethal challenge with ZIKV in mice. **a**, Serum concentration of human mAbs (2 dpi) that was determined by ELISA from mice that were treated prophylactically (d-1) with indicated mAbs (9 μg/mouse) and challenged with ZIKV (d0). Dots show measurements from individual mice (n= 5 to 14 mice per group). Mean ± SD values are shown. **b**, Serum ZIKV titers (2 dpi) from mice treated and challenged as in (a). Dots show measurements from individual mice. Mean ± SD values are shown. **c**, Survival of B6 mice that were treated and challenged as in (a). Overall difference in survival between the groups was estimated using log-rank (Mantel cox) test. P values are indicated. **d**, Low dose (4.5 μg/mouse) prophylaxis of three lead candidate mAbs ZIKV-752, −893, and - 940 (n=5 mice per group), representing each of three competition-binding groups. Mice were treated and challenged as in (a). Overall difference in survival between the groups was estimated using log-rank (Mantel cox) test. P values are indicated. **e**, Low-dose (9 μg/mouse) post-exposure therapeutic efficacy (1 dpi) of three lead candidate mAbs in mice challenged with ZIKV (n= 5 mice per group). Overall difference in survival between the groups was estimated using log-rank (Mantel cox) test. P values are indicated. Data in (a-e) represent single experiments except for ZIKV-752 and −893 in (c) that represent three independent experiments. The mAbs ZIKV-117 and FLU-5J8 or PBS served as controls. Timeline to identify the lead protective mAbs is indicated with a blue arrow in the timeline chart.

Consistent with the reduced viral load in plasma, each of the tested mAbs conferred protection against disease, as shown by reduced and/or delayed mortality compared to the mAb FLU-5J8-treated control group. Three of ten tested mAbs that neutralized both Brazil and Dakar strains (ZIKV-635, −668, and - 893) conferred complete protection (100% survival by 21 dpi), with partial protection conferred by the other two neutralizing mAbs, ZIKV-752 and −940 (**Fig. 5c, Supplemental Table 4**). Protection with a 9 µg dose of ZIKV-893 or −752 was confirmed in three independent experiments. Moreover, similar protection was observed using 2 to 4-fold lower dose of ZIKV-893, −752, or −940 (**Fig. 5d**; **Supplementary Table 4**). The above results demonstrated a high level of efficacy for several of these rapidly identified mAbs against ZIKV when given as prophylaxis.

We next tested therapeutic efficacy after infection was established by administering a low dose of 9 μg of mAb per mouse on 1 dpi. We tested three mAbs ZIKV-893, −752, and −940 because of their broad and potent neutralization of ZIKV, recognition of distinct, non-overlapping epitopes on E (**Fig. 4**), and a high level of protection as prophylaxis (**Supplementary Table 4; Fig. 5a-d**). ZIKV-893 and −940 conferred complete protection that was comparable to the control human mAb ZIKV-117, whereas ZIKV-752 was partially protective (**Fig. 5e; Supplementary Table 4**). A similar level of therapeutic protection was observed using a 2-fold lower dose of ZIKV-752, −893, or −940 mAb (**Supplementary Table 4**).

In an outbreak scenario, an effective rapid public health response incorporating antibody treatments may be limited by the timeline required for protein mAb production in a format that can be safely administered to humans. Nucleic acid-encoded antibody genes offer a potential route to antibody-mediated protection with shorter delivery timelines. As such, and in parallel, we tested therapeutic potency of the twenty lead mAb candidates discovered here with antibody-encoding RNA delivery in mice using a technology that allows *in vivo* mAb expression after intramuscular (i.m.) delivery of an NLC-formulated RNA (J.H.E. et al., paper in preparation). Groups of B6 mice were treated with 40 μg of individual mAb-encoding RNA formulations on day −1 and then challenged with ZIKV Dakar MA on day 0. FLU-5J8- or ZIKV-117-antibody-encoding RNA treated groups served as negative or positive controls, respectively. Human IgG protein was readily detected in the serum of individual mice from most treatment groups (**Fig. 6a**). Five mAbs that were identified to be highly protective with IgG protein treatment (ZIKV-635, −668, −752, - 893, and −940) also mediated a high level of protection with this RNA antibody gene delivery method, as judged by reduced viremia on 2 dpi and survival of mice (**Fig. 6b-c**). These proof-of-concept experiments demonstrate the utility of an antibody-encoding RNA delivery approach for antiviral treatment against ZIKV.

**Fig. 6.**
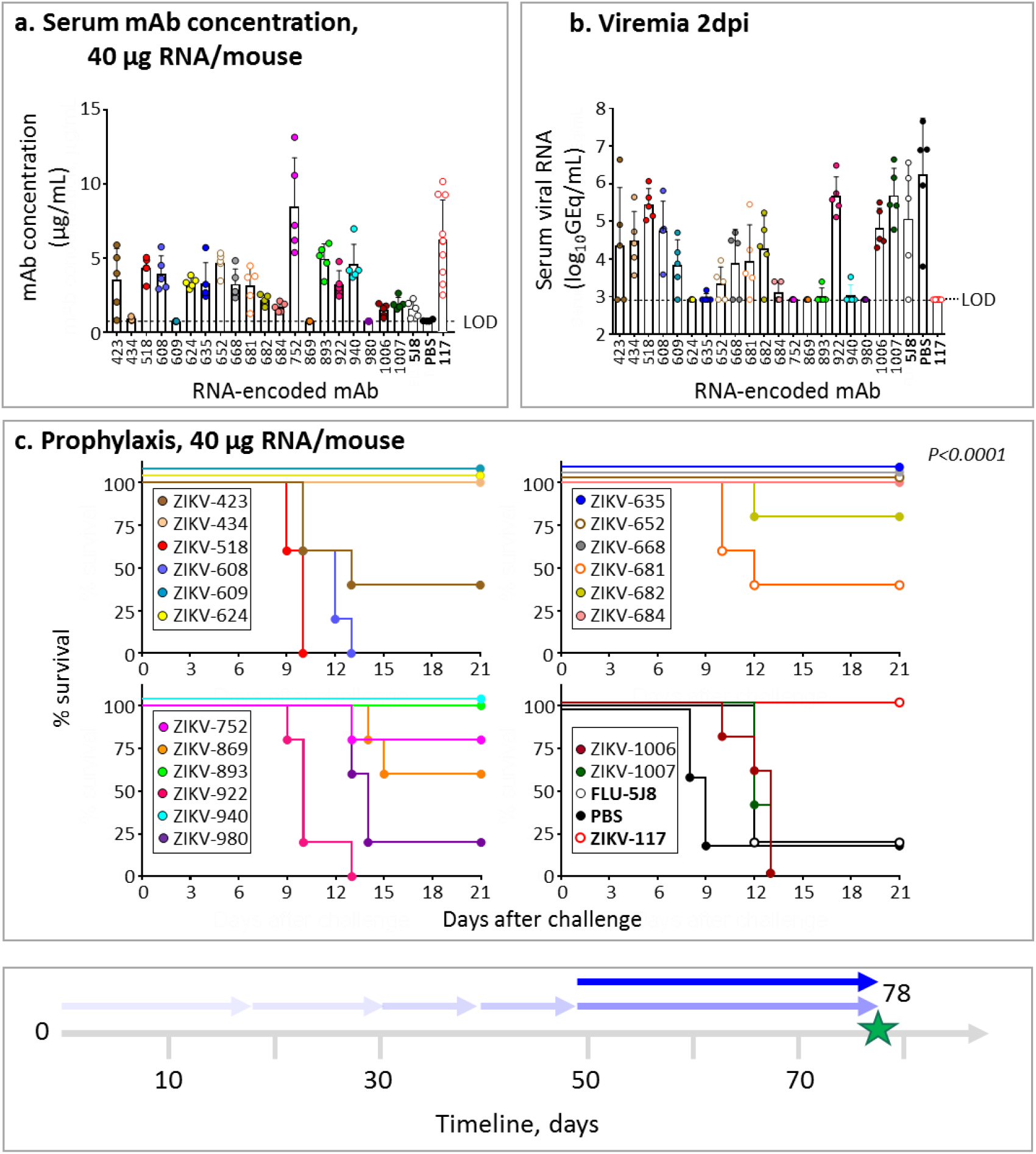
Protection against lethal challenge with ZIKV that was mediated by mAb-encoding RNA formulation delivery in mice. **a**, Serum concentration of human mAbs (2 dpi) from mice that were treated with the indicated mAb-encoding RNA formulations (40 μg/mouse) on day −1 and challenged with ZIKV on day 0 was determined by ELISA. Dots show measurements from individual mice (n=5-10 mice per group). **b**, Serum ZIKV titers (2 dpi) from mice treated and challenged as in (a) was determined by qRT-PCR. Dots show measurements from individual mice. **c**, Survival of mice that were treated and challenged as in (a). Overall difference in survival between the groups was estimated using log-rank (Mantel cox) test. P values are indicated. Data in (a-c) are from one experiment. The mAbs ZIKV-117 and FLU-5J8 served as controls. Timeline to identify the lead protective mAb-encoding RNA formulations is indicated with a blue arrow in the timeline chart.

Together, these results demonstrated the ability to identify rapidly several highly protective neutralizing mAbs against ZIKV and set the stage for evaluation of monotherapy efficacy in the rhesus macaque NHP ZIKV challenge model.

### Modeling a timeline for NHP protection study against ZIKV

At the conclusion of the murine protection studies, we immediately initiated a mAb treatment study of ZIKV infection in NHPs. We produced RNA formulations using a Good Manufacturing Practice (GMP)-compatible process for two lead therapeutic candidate mAbs (ZIKV-752 and −893) in a 200 mg scaled-up process (**Supplementary Fig. 4)** that could be used in NHP studies to enable an Investigational New Drug (IND) application. For the capability demonstration in NHPs, we used our previously described potently neutralizing human mAb ZIKV-117^5^ for the challenge study. We elected to test this antibody in NHPs to maximize the utility of the data, since ZIKV-117 already is on an accelerated clinical development path. Large-scale production and purification of the ZIKV-117 IgG protein used here was not a part of the timeline. The treatment group (n=3 NHPs) received one 10 mg/kg dose of ZIKV-117 intravenously on day −1 and was challenged subcutaneously with 10^3^ PFU of ZIKV strain Brazil the next day. An additional group of NHPs (n=3) that received IgG of irrelevant specificity (sham) served as a control. No detectable viral RNA in plasma and other collected specimens in any of the ZIKV-117-treated NHPs (limit of detection = 2 log_10_ (GEQ/mL)) was observed, whereas each of the control NHPs were positive for infection (**Supplementary Fig. 5a**). ELISA analysis of serum from ZIKV-117 mAb-treated NHPs detected ∼50 μg/mL of circulating ZIKV-117 the day after the treatment (**Supplementary Fig. 5b**). Together, this timeline modeling study allowed us to define the efficacy of mAb monotherapy against ZIKV infection in NHPs in only nine days.

## Discussion

We describe an integrated technology platform pipeline modeling deployment of the rapid response for discovery of antiviral antibodies. Recently, several laboratories have demonstrated the feasibility of rapid identification of potently antiviral mAbs^12^. The shortest reported time frame of which we are aware to date is about 4 months, which was reported for isolation of neutralizing mAb LCA60 against MERS-CoV, a process that included mAb identification, functional screening, mouse protection study and development of a stable cell line for mAb production^34^. Our study demonstrated that using a pipeline of integrated technologies, human antiviral human mAb discovery and therapeutic potency verification (including mAb RNA delivery and proof-of principle NHP protection study), could be accomplished in 78 days, 12 days shorter than our 90 day goal. It should be noted that current single-cell antibody gene cloning techniques permit the rapid generation of antigen-reactive mAbs in 2 weeks or even less^35,36^. However, isolation of mAbs that would be suitable for clinical development (*e.g.* mAbs with exceptional *in vitro* and *in vivo* potency), remains a bottleneck of antibody discovery process^12^. We demonstrated high performance of an integrated technology platform pipeline for rapid discovery of potent human antiviral mAbs. A comparison to historical studies with previously reported human mAbs against ZIKV^6,7,18,23^ qualifies the newly identified broadly-neutralizing mAbs ZIKV-752, −893, and −940 as promising therapeutic candidates of high potency against ZIKV. In addition, these mAbs recognize diverse epitopes, which has relevance for therapeutic cocktails with complementary specificity and activity. Although a conventional mouse challenge model to assess protection against ZIKV uses long timelines (typically 16-21 days per study), early examination of viremia and a mAb dose-de-escalation approach could be used to identify highly protective mAbs in a short timeframe required by rapid response programs. Together, these results suggest that development and use of mAbs as alternative antiviral therapeutics could be practical when compared to conventional antiviral countermeasures, such as vaccines or small molecule antiviral drugs. Continuing improvement of instrument platforms for high-throughput single cell analysis (DeKosky et al., 2013; Setliff et al., 2019) warrants feasibility of rapid antibody discovery for emerging viruses in the future.

One of the remaining challenges for widespread application of antiviral antibody therapies is the potentially higher cost associated with production for intravenous administration. An alternative to passive IgG protein administration is nucleic acid delivery of genes encoding antibodies using cDNA or mRNA. A prior study showed that DNA-encoded intramuscular mAb delivery protected NHPs from ZIKV challenge after three sequential DNA administrations^37^. Another study reported NHP protection against chikungunya virus infection after intravenous infusion of a lipid nanoparticle-encapsulated mRNA (LNP) encoding antiviral mAb^38^, and this formulation was used in a first-in-man clinical trial of mRNA-encoded antibodies in 2019 (ClinicalTrials.gov Identifier: NCT03829384). The NLC formulation that we used in this study may have an advantage over lipid nanoparticle encapsulated RNA formulation^39^, in that the RNA is externally bound to the nanostructured lipid carrier, as opposed to the internally placed RNA of the LNP formulation. This approach could allow for stockpiling of the stable nanoparticles without cold chain manufacturing. Thus, final formulation could be performed as needed once the desired mRNA encoding an antibody is identified and produced. Formulated RNA allows high level of mAb expression after intramuscular injection *in vivo* in mice (J.H.E. et al., paper in preparation). The extent to which these findings translate to NHPs and humans remains unclear, and this aspect of the approach must be determined in future studies. Nonetheless, our study provides a roadmap for the approach to large-scale accelerated discovery of potent antiviral human mAb therapeutics against known viral antigens and sets the stage for pre-clinical evaluation of mAbs using RNA delivery platforms.

## Methods

### Research subjects

We studied seven subjects in the United States with previous or recent ZIKV infection (**Supplementary Table 1)**. The studies were approved by the Institutional Review Board of Vanderbilt University Medical Center; samples were obtained after written informed consent was obtained by the Vanderbilt Clinical Trials Center. Subjects were infected during the 2015-2016 outbreak of an Asian lineage strain, following exposure in Brazil, Nicaragua, Puerto Rico, Dominican Republic, Guatemala or Haiti.

### Animals

Mouse challenge studies were approved by the Washington University School of Medicine (Assurance number A3381-01) Institutional Animal Care and Use Committee. The facility where animal studies were conducted is accredited by the Association for Assessment and Accreditation of Laboratory Animal Care, International and follows guidelines set forth by the Guide for the Care and Use of Laboratory Animals, National Research Council, 2011. Blood was obtained as approved by submandibular vein bleeding. Virus inoculations were performed under anesthesia that was induced and maintained with ketamine hydrochloride and xylazine, and all efforts were made to minimize animal suffering. Mice were non-specifically and blindly distributed into respective groups.

The NHP research studies adhered to principles stated in the eighth edition of the Guide for the Care and Use of Laboratory Animals. The facility where this research was conducted [(Alpha Genesis Incorporated, Yemassee, SC)] is fully accredited by the Association for Assessment and Accreditation of Laboratory Animal Care International and has an approved Office of Laboratory Animal Welfare Assurance (A3645-01).

### Cell culture

Vero cells (American Type Culture Collection) were maintained at 37°C in 5% CO_2_ in Dulbecco’s minimal essential medium (DMEM) containing 10% (vol/vol) heat-inactivated fetal bovine serum (FBS) and 1 mM sodium pyruvate. All cell lines were tested and found negative for mycoplasma contamination.

### Viruses

Previously described human isolates of ZIKV Brazil, Paraiba 01/2015 (GenBank: KX280026.1)^21,40^, and ZIKV Dakar 41525, Senegal 1984 (GenBank: KU955591)^20^ were used for *in vitro* studies. A previously described mouse-adapted isolate of ZIKV Dakar 41525^20^ ZIKV Dakar MA (GenBank: MG758786.1), was used for both *in vitro* and *in vivo* studies. For NHP studies a human ZIKV strain from Brazil (Brazil 2015) was used^41,42^.

### Virus stock production

To identify cells suitable for production of high-titer ZIKV stocks in an antigen-independent manner, we used qRT-PCR to rapidly detect ZIKV viral RNA in a panel of immortalized cell lines including those commonly used for virus propagation (Vero, BHK, JEG-3, HeLa, HEK-293T, U20S, A549, Huh7, Huh7.5, EA.Hy926, and Hap-1 cell lines). For subsequent ZIKV production studies, we chose Vero cells due to their well-characterized use in virus production. To generate stocks of ZIKV for subsequent *in vitro* and *in vivo* studies, Vero cells plated in T175 flasks were inoculated with a seed stock and cell culture supernatant collected 66 to 72 hrs after inoculation and titrated. Virus stocks were then titrated by focus forming assay (FFA), aliquoted, and stored at −80°C until use. To identify conditions suitable for large-scale production of ZIKV stocks in Vero cells grown in roller bottles, we examined ZIKV titers in various serum concentrations ranged from 0.5 to 5% of FBS in cell culture supernatant collected 24 to 72 hrs after inoculation. We observed that even 0.5% fetal bovine serum was sufficient to generate high-titer ZIKV stocks by 72 hrs after virus inoculation. To identify conditions for concentration and inactivation of virus stocks, we determined the infectivity of ZIKV stocks by FFA following ultracentrifugation as previously described^43^ and following inactivation with hydrogen peroxide in an attempt to preserve immunogenicity as described previously^44^. We observed increased ZIKV titers in the virus pellet following ultra-centrifugation and could not detect infectious ZIKV following treatment with 3% hydrogen peroxide for 1 h at ambient temperature and 0.024 U/μL of catalase for 10 min at ambient temperature (to inactivate hydrogen peroxide).

### Human subjects selection and target-specific memory B cells isolation

B cell responses to ZIKV in PBMCs from a cohort of 11 subjects with previous exposure to the ZIKV Asian lineage were assessed to identify subjects with the highest response. The frequency of ZIKV-specific B cells was enumerated from frozen PBMCs using an African ZIKV lineage soluble recombinant E protein (Meridian Bioscience), the antigen that was used previously to identify potent ZIKV mAbs^5^. Briefly, B cells were purified magnetically (STEMCELL Technologies) and stained with anti-CD19, - IgD, -IgM, and -IgA phenotyping antibodies (BD Biosciences) and biotinylated E protein. 4′,6-diamidino-2-phenylindole (DAPI) was used as a viability dye to discriminate dead cells. Antigen-labeled class-switched memory B cell-E complexes (CD19^+^IgM^-^IgD^-^IgA^-^ZIKV E^+^DAPI^-^) were detected with allophycocyanin (APC)-labeled streptavidin conjugate and quantified using an iQue Plus Screener flow cytometer (IntelliCyt Corp). After identification of the seven subjects with the highest B cell response against ZIKV, target-specific memory B cells were isolated by FACS using an SH800 cell sorter (Sony) from pooled PBMCs of these subjects, after labeling of B cells with biotinylated E protein (sub-panel 1).

Although sorting with a biotinylated E antigen was used successfully for isolation of human neutralizing ZIKV mAbs, this approach might fail to identify mAbs whose antigenic sites altered by the biotin labeling of the target protein. To overcome this limitation, we also used alternate approaches. The second approach used binding to soluble intact E protein antigen, but included a labeled antibody detected at the E fusion loop (FL) (indirect label method, sub-panel 2). This approach circumvented any binding issues stemming from biotin-labeling as well as excluded the subset of B cells that encode FL-specific mAbs, which dominate in the response to ZIKV but typically poorly neutralize the virus^5^. Third, all labeled B cells (as in the mAb sub-panel 2) and a subset of high-affinity B cells (high E labeling clones, mAb sub-panel 3) were sorted separately. Overall, from > 5 x 10^8^ PBMCs, > 5,000 E-specific B cells were sorted and subjected to further analysis. Two methods were implemented for the preparation of sorted B cells for sequencing. Approximately 800 sorted cells were subjected to direct sequencing immediately after FACS. The remaining cells were expanded in culture for eight days in the presence of irradiated 3T3 feeder cells that were engineered to express human CD40L, IL-21, and BAFF as described previously^45^. The expanded lymphoblastoid cell lines (LCLs) secreted high levels of E-specific mAbs, as confirmed by ELISA from culture supernatants. Approximately 40,000 expanded LCLs were sequenced using the Chromium sequencing method (10x Genomics).

### Generation of single cell antibody variable genes profiling libraries

The Chromium Single Cell V(D)J workflow with B-cell only enrichment option was chosen for generating linked heavy-chain and light-chain antibody profiling libraries. Approximately 800 directly sorted E-specific B cells were split evenly into two replicates and separately added to 50 μL of RT Reagent Mix, 5.9 μL of Poly-dt RT Primer, 2.4 μL of Additive A and 10 μL of RT Enzyme Mix B to complete the Reaction Mix as per the vendor’s protocol, which then was loaded directly onto a Chromium chip (10x Genomics). Similarly, for the remaining sorted cells that were bulk expanded, approximately 40,000 cells from two separate sorting approaches were split evenly across six reactions and separately processed as described above before loading onto a Chromium chip. The libraries were prepared following the User Guide for Chromium Single Cell V(D)J Reagents kits (CG000086_REV C).

### NGS sequencing

Chromium Single Cell V(D)J B-Cell enriched libraries were quantified, normalized and sequenced according to the User Guide for Chromium Single Cell V(D)J Reagents kits (CG000086_REV C). The two enriched libraries from direct flow cytometric cell sorting were combined into one library pool and sequenced on a NovaSeq sequencer (Illumina) with a NovaSeq 6000 S1 Reagent Kit (300 cycles) (Illumina). The six enriched libraries from bulk expansion were combined into one library pool and sequenced on a NovaSeq sequencer with a NovaSeq 6000 S4 Reagent Kit (300 cycles (Illumina). All enriched V(D)J libraries were targeted for sequencing depth of at least 5,000 raw read pairs per cell. Following sequencing, all samples were demultiplexed and processed through the 10x Genomics Cell Ranger software (version 2.1.1) as detailed below.

### Bioinformatics analysis

The down-selection to identify lead candidates for expression was carried out in two phases. In the first phase, all paired antibody heavy and light chain variable gene cDNA nucleotide sequences obtained that contained a single heavy and light chain sequence were processed using our Python-based antibody variable gene analysis tool (PyIR; https://github.com/crowelab/PyIR) (C.S. et al., paper in preparation). We considered heavy and light chain encoding gene pairs productive and retained them for additional downstream processing if they met the following criteria: 1) did not contain a stop codon, 2) had an intact CDR3 and 3) contained an in-frame junctional region. The second phase of processing grouped all productive heavy chain nucleotide sequences based on their V3J clonotype (defined by identical inferred V and J gene assignments and identical amino acid sequences of the CDR3)^46^. All heavy chain nucleotide sequences from individual sub-panels were grouped by their heavy chain V3J clonotype. The somatic variants belonging to each heavy chain V3J clonotype grouping then were rank-ordered based on the degree of somatic mutation, and only the most mutated heavy chain sequence was retained for downstream expression and characterization. Any somatic variant that was not designated as an IgG isotype (based on the sequence and assignment using the 10x Genomics Cellranger V(D)J software [version 2.1.1]) was removed from consideration. All heavy and light chain nucleotide sequences were translated, and redundant entries removed to avoid expressing identical mAbs. The identities of antibody variable gene segments, CDRs, and mutations from inferred germline gene segments were determined by alignment using the ImMunoGeneTics database^47^.

### Antibody gene synthesis

Sequences of selected mAbs were synthesized using a rapid high-throughput cDNA synthesis platform (Twist Bioscience) and subsequently cloned into an IgG1 monocistronic expression vector (designated as pTwist) for mammalian cell culture mAb secretion. This vector contains an enhanced 2A sequence and GSG linker that allows simultaneous expression of mAb heavy and light chain genes from a single construct upon transfection ^25^.

### MAb production and purification

For high-throughput production of recombinant mAbs, we adopted approaches designated as “micro-scale”. For mAbs expression, we performed micro-scale transfection (∼1 mL per antibody) of Chinese hamster ovary (CHO) cell cultures using the Gibco™ ExpiCHO™ Expression System and a protocol for deep 96-well blocks (ThermoFisher Scientific). Briefly, synthesized antibody-encoding lyophilized DNA (∼2 μg per transfection) was reconstituted in OptiPro serum free medium (OptiPro SFM), incubated with ExpiFectamine CHO Reagent and added to 800 µL of ExpiCHO cell cultures into 96 deep well blocks using a ViaFlo 384 liquid handler (Integra Biosciences). Plates were incubated on an orbital shaker at 1,000 rpm with 3 mm orbital diameter at 37°C in 8% CO_2_. The next day after transfection, ExpiFectamine CHO Enhancer and ExpiCHO Feed reagents were added to the cells, followed by 4 days incubation for a total of 5 days at 37°C in 8% CO_2_. Culture supernatants were harvested after centrifugation of blocks at 450 x *g* for 5 min and stored at 4°C until use. For high-throughput micro-scale mAbs purification, we used fritted deep well plates containing 25 μL of settled protein G resin (GE Healthcare Life Sciences) per well. Clarified culture supernatants were incubated with protein G resin for mAb capturing, washed with PBS using a 96-well plate manifold base (Qiagen) connected to the vacuum, and eluted into 96-well PCR plates using 86 μL of 0.1 M glycine-HCL buffer pH 2.7. After neutralization with 14 μL of 1 M Tris-HCl pH 8.0, purified mAbs were buffer exchanged into PBS using Zeba™ Spin Desalting Plates (ThermoFisher Scientific) and stored at 4°C until use.

For scaling up of mAb production for *in vivo* mouse studies, the transfection was performed in 125 mL Erlenmeyer vented cap flasks (Corning) containing 35 mL of ExpiCHO cells following the manufacturer’s protocol. Antibodies were purified from filtered culture supernatants by fast protein liquid chromatography (FPLC) on an ÄKTA Pure instrument using a HiTrap MabSelect Sure column (GE Healthcare Life Sciences). Purified mAbs were buffer exchanged into PBS, filtered using sterile 0.45-μm pore size filter devices (Millipore), concentrated, and stored in aliquots at 4°C until use. To quantify purified mAbs, absorption at 280 nm (A280) was measured using NanoDrop (ThermoFisher Scientific), and mAb concentration was calculated using IgG sample type setting on NanoDrop that uses a typical for IgG molar extinction coefficient equal to 210,000 M^-1^cm^-1^.

### Antibody-encoding RNA production

cDNAs encoding the 20 lead mAb candidates were amplified from the mammalian expression pTwist vector using universal primer sets and cloned into plasmids encoding Venezuelan equine encephalitis virus replicon (strain TC-83) under the control of a T7 promoter (pT7-VEE-Rep)^32^. To generate linear templates for RNA transcription for RNA *in vitro* transcription and capping at midi scale (500 μg), plasmid DNA was cut by restriction digest using NotI or BspQI enzymes (New England Biolabs), respectively, and purified using phenol chloroform extraction and sodium acetate precipitation. RNA was transcribed using T7 Polymerase, RNAse inhibitor, Pyrophosphatase enzymes (Aldevron) and reaction buffer. RNA transcripts were capped with vaccinia virus capping enzyme using GTP and S-adenosyl-methionine (Aldevron) as substrates to create a cap-0 structure. RNA was purified using lithium chloride precipitation. Large-scale capability demonstration RNA production was performed using a 200 mg scaled-up process and purified via column on a Capto Core 700 instrument (GE Healthcare Life Sciences) and tangential flow filtration.

### RNA formulation production

The nanostructured lipid carrier (NLC) nanoparticle formulation was manufactured as previously described ^32^. The oil phase is composed of squalene (the liquid-phase of the oil core), glyceryl trimyristate (Dynasan® 114) (the solid-phase of the oil core), a non-ionic sorbitan ester surfactant [sorbitan monostearate (Span® 60)], and the cationic lipid DOTAP (N-[1-(2,3-dioleoyloxy)propyl]-N,N,N-trimethylammonium chloride). The aqueous phase is a 10 mM sodium citrate trihydrate buffer containing the non-ionic PEGylated surfactant Tween® 80. Separately, the two phases were heated and equilibrated to 60°C in a bath sonicator. Following complete dissolution of the solid components, the oil and aqueous phases were mixed at 5,000 rpm in a high-speed laboratory emulsifier (Silverson Machines, Inc.) to produce a crude mixture containing micron-sized oil droplets. Further particle size reduction was achieved by high-shear homogenization in a M-110P microfluidizer (Microfluidics, Corp.). The colloid mixtures were processed at 30,000 psi for five discrete microfluidization passes. The final pH was adjusted to between 6.5 and 6.8. Formulations were filtered with a 0.2 µm polyethersulfone membrane syringe filter and stored at 2 to 8°C. RNA was complexed with NLC at a nitrogen-to-phosphate ratio of 5, diluting NLC in 10 mM citrate buffer and RNA in a 20% sucrose solution and complexing on ice for 30 min prior to use.

### ELISA binding screening assays

Wells of 96-well microtiter plates were coated with purified recombinant ZIKV E protein (Meridian Bioscience) at 4°C overnight. Plates were blocked with 2% non-fat dry milk and 2% normal goat serum in DPBS containing 0.05% Tween-20 (DPBS-T) for 1 hr. For mAb screening assays, CHO cell culture supernatants or purified mAbs were diluted 1:20 in blocking buffer, added to the wells, and incubated for 1 hr at ambient temperature. The bound antibodies were detected using goat anti-human IgG conjugated with HRP (horseradish peroxidase) (Southern Biotech) and TMB (3,3µ,5,5µ-tetramethylbenzidine) substrate (ThermoFisher Scientific). Color development was monitored, 1N hydrochloric acid was added to stop the reaction, and the absorbance was measured at 450 nm using a spectrophotometer (Biotek). For dose-response assays, serial dilutions of purified mAbs were applied to the wells in triplicate or quadruplicate, and mAb binding was detected as detailed above.

### Focus reduction neutralization test

Serial dilutions of mAbs were incubated with 10^2^ FFU of different ZIKV strains (Dakar or Brazil) for 1 h at 37°C. The mAb–virus complexes were added to Vero cell monolayers in 96-well plates for 90 min at 37°C. Subsequently, cells were overlaid with 1% (w/v) methylcellulose in Minimum Essential Medium (MEM) supplemented with 4% heat-inactivated FBS. Plates were fixed 40 hrs later with 1% paraformaldehyde (PFA) in PBS for 1 h at room temperature. The plates were incubated sequentially with 500 ng/mL previously described mouse anti-ZIKV mAb ZV-16^5^ and horseradish-peroxidase (HRP)-conjugated goat anti-mouse IgG in PBS supplemented with 0.1% (w/v) saponin (Sigma) and 0.1% bovine serum albumin (BSA). ZIKV-infected cell foci were visualized using TrueBlue peroxidase substrate (KPL) and quantitated on an ImmunoSpot 5.0.37 Macro Analyzer (Cellular Technologies).

### High-throughput mAbs quantification

High-throughput quantification of micro-scale produced mAbs was performed from CHO culture supernatants or micro-scale purified mAbs in a 96-well plate format using the Cy-Clone Plus Kit and an iQue Plus Screener flow cytometer (IntelliCyt Corp), according to the vendor’s protocol. Purified mAbs were assessed at a single dilution (1:10 final, using 2 μL of purified mAb per reaction), and a control human IgG solution with known concentration was used to generate a calibration curve. Data were analyzed using ForeCyt software version 6.2 (IntelliCyt Corp).

### Real-time cell analysis assay (RTCA)

To screen for neutralizing activity in the panel of recombinantly expressed mAbs, we used a high-throughput and quantitative real-time cell analysis (RTCA) assay and xCelligence Analyzer (ACEA Biosciences Inc.) that assesses kinetic changes in cell physiology, including virus-induced cytopathic effect (CPE). Fifty (50) μL of cell culture medium (DMEM supplemented with 2% FBS) was added to each well of a 96-well E-plate using a ViaFlo384 liquid handler (Integra Biosciences) to obtain background reading. Eighteen thousand (18,000) Vero-E6 cells in 50 μL of cell culture medium were seeded per each well, and the plate was placed on the analyzer. Measurements were taken automatically every 15 min, and the sensograms were visualized using RTCA software version 2.1.0 (ACEA Biosciences Inc). For a screening neutralization assay, the virus (0.5 MOI, ∼9,000 PFU per well) was mixed with a 1:10 dilution of CHO supernatants obtained from micro-scale expression experiments, or a 1:25 dilution of micro-scale purified Abs in a total volume of 50 μL using DMEM supplemented with 2% FBS as a diluent and incubated for 1 h at 37°C in 5% CO_2_. At ∼12 hrs after seeding the cells, the virus-mAb mixtures were added in replicates to the cells in 96-well E-plates. Wells containing virus only (in the absence of mAb) and wells containing only Vero cells in medium were included as controls. Plates were measured continuously (every 15 minutes) for 48 to 72 hrs to assess virus neutralization. A mAb was considered as neutralizing if it partially or fully inhibited ZIKV-induced CPE. For mAb potency ranking experiments, individual mAbs identified as fully neutralizing from the screening study were assessed at six five-fold dilutions starting from 1 μg/mL in replicates after normalization of mAb concentrations. IC_50_ values were estimated as cellular index (CI) change over time, using non-linear fit with variable slope analysis performed in the RTCA software version 2.1.0 (ACEA Biosciences Inc.). The potency ranking study was repeated using midi-scale purified mAbs to confirm the activity and to assure the quality of antibody preparations before performing *in vivo* protection studies in mice.

### Antibody-mediated cellular phagocytosis by human monocytes (ADCP)

Recombinant soluble ZIKV E protein was biotinylated and coupled to AF488 Neutravidin beads (Life Technologies). Microscale-purified mAbs were tested at a single 1:10 dilution; concentrations were not normalized. Antibodies were diluted in cell culture medium and incubated with beads for 2 h at 37°C. THP-1 monocytes (ATCC) were added at 2.5 x 10^4^ cells per well and incubated for 18 h at 37°C. Cells were fixed with 4% PFA and analyzed on a BD LSRII flow cytometer, and a phagocytic score was determined using the percentage of AF488^+^ cells and the MFI of the AF488^+^ cells. The previously identified human mAb ZIKV-117 was used as a positive control, and human mAb FLU-5J8 was used as a negative control.

### Antibody-mediated neutrophil phagocytosis (ADNP)

Recombinant soluble ZIKV E protein was biotinylated and coupled to AF488 Neutravidin beads (Life Technologies). Microscale-purified mAbs were tested at a single 1:10 dilution; concentrations were not normalized. Antibodies were diluted in cell culture medium and incubated with beads for 2 hrs at 37°C. White blood cells were isolated from the peripheral blood of subjects by lysis of red blood cells, followed by three washes with PBS. Cells were added at a concentration of 5.0 x 10^4^ cells/well and incubated for 1 hr at 37°C. Cells were stained with CD66b (Pacific Blue, Clone G10F5; BioLegend), CD3 (AF00, Clone UCHT1; BD Biosciences), and CD14 (APC-Cy7, Clone MφP9; BD Biosciences), and fixed with 4% PFA, and analyzed by flow cytometry on a BD LSR II flow cytometer. Neutrophils were defined as SSC-A^high^ CD66b+, CD3-, CD14-. A phagocytic score was determined using the percentage of AF488^+^ cells and the MFI of the AF488^+^ cells. The previously identified mAb ZIKV-117 was used as a positive control, and mAb FLU-5J8 was used as a negative control.

### Antibody-mediated complement deposition (ADCD)

Recombinant soluble ZIKV E protein was biotinylated and coupled to red fluorescent Neutravidin beads (Life Technologies). Antibodies were diluted to 5 μg/mL in RPMI-1640, and incubated with GP-coated beads for 2 h at 37°C. Freshly reconstituted guinea pig complement (Cedarlane Labs) was diluted in veronal buffer with 0.1% fish gelatin (Boston Bioproducts), added to the antibody-bead complexes, and incubated for 20 min at 37°C. Beads were washed twice with PBS containing 15 mM EDTA, and stained with an anti-guinea pig C3 antibody conjugated to FITC (MP Biomedicals) for 15 min at ambient temperature. Beads were washed twice more with PBS, and C3 deposition onto beads was analyzed on a BD LSRII flow cytometer and the median fluorescence intensity of the FITC+ of all beads was measured.

### Competition-binding analysis and epitope mapping

For the competition-binding assay, wells of 384-well microtiter plates were coated with purified recombinant ZIKV E protein at 4°C overnight. Plates were blocked with 2% BSA in DPBS containing 0.05% Tween-20 (DPBS-T) for 1 hr. Purified unlabeled mAbs were diluted in blocking buffer to 5 μg/mL, added to the wells (25 μL/well), and incubated for 1 h at ambient temperature. Biotinylated previously characterized ZIKV E-specific human mAbs ZIKV-117, ZIKV-116, and ZIKV-88 were added to indicated wells at 5 μg/mL in a 2.5 μL/well volume (final ∼0.5 μg/mL concentration of biotinylated mAb) without washing of unlabeled antibody and then incubated for 1 h at ambient temperature. Plates were washed, and bound antibodies were detected using HRP-conjugated avidin (Sigma) and TMB substrate. The signal obtained for binding of the biotin-labeled reference antibody in the presence of the unlabeled tested antibody was expressed as a percentage of the binding of the reference antibody alone after subtracting the background signal. Tested mAbs were considered competing if their presence reduced the reference antibody binding to less than 41% of its maximal binding and non-competing if the signal was greater than 71%. A level of 40–70% was considered intermediate competition.

Epitope mapping was performed by shotgun mutagenesis essentially as described previously^48^. A ZIKV prM/E protein expression construct (based on ZIKV strain SPH2015) was subjected to high-throughput alanine scanning mutagenesis to generate a comprehensive mutation library. Each residue within prM/E was changed to alanine, with alanine codons mutated to serine. In total, 672 ZIKV prM/E mutants were generated (100% coverage), sequence confirmed, and arrayed into 384-well plates. Each ZIKV prM/E mutant was transfected into HEK-293T cells and allowed to express for 22 hrs. Cells were fixed in 4% (v/v) paraformaldehyde (Electron Microscopy Sciences), and permeabilized with 0.1% (w/v) saponin (Sigma-Aldrich) in PBS plus calcium and magnesium (PBS++). Cells were incubated with purified mAbs diluted in PBS++, 10% normal goat serum (Sigma), and 0.1% saponin. Primary antibody screening concentrations were determined using an independent immunofluorescence titration curve against wild-type ZIKV prM/E to ensure that signals were within the linear range of detection. Antibodies were detected using 3.75 μg/mL of AlexaFluor488-conjugated secondary antibody (Jackson ImmunoResearch Laboratories) in 10% normal goat serum with 0.1% saponin. Cells were washed three times with PBS++/0.1% saponin followed by two washes in PBS. Mean cellular fluorescence was detected using a high-throughput flow cytometer (HTFC, Intellicyt Corp.). Antibody reactivity against each mutant prM/E clone was calculated relative to wild-type prM/E protein reactivity by subtracting the signal from mock-transfected controls and normalizing to the signal from wild-type prM/E-transfected controls. Mutations within clones were identified as critical to the mAb epitope if they did not support reactivity of the test MAb, but supported reactivity of other ZIKV antibodies. This counter-screen strategy facilitates the exclusion of prM/E mutants that are locally misfolded or have an expression defect.

### Detection of circulating human mAbs in mouse serum

The amount of human mAb in serum was detected using a capture ELISA with a standard curve of recombinant anti-ZIKV mAb ZIKV-117 or an IgG1 isotype-matched control anti-influenza mAb, FLU-5J8. Briefly, plates were coated with 2 µg/mL of goat anti-human kappa or goat anti-human lambda antibodies cross-absorbed against mouse IgG (Southern Biotech) at 4°C overnight. Plates were blocked using 2% BSA in PBS for 1 h at 37°C and then incubated for 1 h at 4°C with serial dilutions of heat-inactivated mouse serum in parallel with a serial dilution of a known quantity of ZIKV-117 hIgG1 protein. After washing, bound antibody was detected using HRP-conjugated goat anti-human Fc multiple species cross-absorbed (Southern Biotech) for 1 h at 4°C. Plates were developed using TMB substrate (Thermo Fisher Scientific), and the reaction was stopped with H_2_SO_4_. ELISA plates were read using a TriBar LB941 plate reader (Berthold Technologies). The optical density values from the known quantity of ZIKV-117 hIgG1 were fitted to a standard curve and compared to the optical density values of serum to determine the concentration of ZIKV-117 hIgG1.

### ZIKV titer measurements

Blood was collected from ZIKV-infected mice at various time points, allowed to clot at ambient temperature, and serum was separated by centrifugation. Viral RNA was isolated using the 96-well Viral RNA kit (Epigenetics), as described by the manufacturer. ZIKV RNA levels were determined by TaqMan one-step qRT-PCR, as described previously^49^. Alternatively, ZIKV Dakar MA and Brazil strain titers were determined by FFA on Vero cell monolayer cultures, as previously described^50^.

### Mouse challenge experiments

Wild-type C57BL/6J mice (4 weeks of age) were purchased from Jackson Laboratory. For studies involving ZIKV challenge, mice were treated with 2 mg of an Ifnar1-blocking antibody (MAR1-5A3, Leinco Technologies) by i.p. injection one-day before virus inoculation. For prophylaxis experiments, mice were treated i.p. one day before virus challenge with purified individual mAbs and monitored for 21 days for survival. For therapeutic protection experiments, mice were treated i.p. one day after virus challenge with varying defined doses of purified individual Abs and monitored for 21 days for survival. For antibody-encoding RNA delivery studies, mice were inoculated with 40 µg of individual RNA by an i.m. route (into each quadricep and hamstring muscle groups) one day before virus challenge, with four injections of 50 μL of RNA/NLC complex per injection. For ZIKV infections, mice were inoculated by a subcutaneous (via footpad) route with 10^3^ FFU of ZIKV Dakar MA in a volume of 30 μL of PBS. Antibody ZIKV-117 was used as a positive control, and mAb FLU-5J8 or PBS were used as a negative control.

### NHP challenge study

Six healthy adult rhesus macaques (*Macaca mulatta*) of Chinese origin (5 to 15 kg body weight) were studied. Animals were allocated randomly to two treatment groups (n=3 per group) and two control (“sham”-treated) groups (n=3 per group). All animals were inoculated by subcutaneous (s.q). route with a target dose of 10^6^ viral particles (approx. 10^3^ PFU) of ZIKV Brazil prior to mAb infusions. The macaques in the treatment group received 10 mg/kg of ZIKV-117 mAb by intravenous injection one day prior ZIKV challenge. The control sham-treated animals received 10 mg/kg of mAb PGT121 that is specific to gp120 protein antigen of HIV-1 one day prior ZIKV challenge. All animals were given physical exams, and blood was collected at the time of ZIKV inoculation and at indicated times after ZIKV inoculation. In addition, all animals were monitored daily with an internal scoring protocol approved by the Institutional Animal Care and Use Committee. These studies were not blinded.

### Detection of virus load in NHPs by qRT-PCR analysis

Titration of virus in indicated specimens was performed by qRT-PCR analysis as previously described^41,42^. Viral RNA was isolated from plasma and other tested specimens with a QIAcube HT (QIAGEN, Germany). A QIAcube 96 Cador Pathogen kit or RNeasy 96 QIAcube HT kit were used for RNA extraction. A cDNA of the wild-type BeH815744 Cap gene was cloned into the pcDNA3.1 expression plasmid, then transcribed *in vitro* to RNA using the AmpliCap-Max T7 High Yield Message Maker Kit (Cellscript). RNA quality was assessed by the Beth Israel Deaconess Medical Center molecular core facility. Ten-fold dilutions of the RNA were prepared for standards and reverse transcribed to cDNA. Primers were synthesized by Integrated DNA Technologies (Coralville), and probes were obtained from Biosearch Technologies (Petaluma). Viral loads were calculated as virus particles (VP) per mL, and the assay sensitivity was 100 copies/mL.

### Detection of circulating human mAbs in NHP serum

A modified protocol using a commercially available human anti-zika virus Envelope protein (ZIKV-Env) IgG kit (Alpha Diagnostics International) was used to quantify ZIKV-117 mAb levels in NHP serum samples. In brief, plate wells were equilibrated several times with kit NSB wash buffer and 100 µL of standard calibrators, and NHP serum dilutions were prepared in kit sample diluent buffer plated in duplicate and incubated for 1 h at ambient temperature. Plates were washed and NHP IgG pre-adsorbed HRP-conjugated anti-human secondary antibodies (Novus Biological) diluted to 1:2,000 in kit sample diluent buffer was added at 100 µL/well and incubated for 1 h. Plates were washed and developed for two minutes using the kit TMB solution, stopped by addition of kit stop solution and analyzed at 450 nm/550 nm on a Versamax microplate plate reader (Molecular Devices) using Softmax Pro 6.5.1 software. A four-parameter logistic (4-PL) standard curve was generated using Prism v8.0 software (GraphPad) for the standard calibrators, and the concentrations of the unknown samples were interpolated from the linear portion of the curve.

### Quantification and statistical analysis

The descriptive statistics mean ± SEM or mean ± SD were determined for continuous variables as noted. Survival curves were estimated using the Kaplan-Meier method, and an overall difference between groups was estimated using the two-sided log rank test (Mantel-Cox) with subjects right censored, if they survived until the end of the study. In FRNT neutralization assays, IC_50_ values were calculated after log transformation of antibody concentrations using a 3-parameter nonlinear fit analysis. In the RTCA neutralization assays, IC_50_ values were estimated as cellular index change over time using non-linear fit with variable slope analysis determined in the RTCA software version 2.1.0 (ACEA Biosciences Inc.). Technical and biological replicates are described in the figure legends. Statistical analyses were performed using Prism v7.2 (GraphPad).

## Acknowledgements

We thank M. Mayo for assistance with acquisition of the human survivor samples and coordination across study sites, J. Govero for assistance with RNA protection experiments, A. Jones and K. Beeri for assistance and coordination of NGS sequencing timelines, J.C. Slaughter, M. Goff and R. Troseth for assistance with data analysis, STEMCELL Technologies and ACEA Biosciences for providing resources. This study was supported by Defense Advanced Research Projects Agency (DARPA) grant HR0011-18-2-0001 and HHS contract HHSN272201400058C (to J.E.C. and B.J.D.). The content is solely the responsibility of the authors and does not necessarily represent the official views of the DARPA.

## Contributions

P.G., R.G.B., J.H.E, R.N., C.S., T. J. S., E.D., B.J.D., T.J.S., L.T., G.A., S.G. R., N.V.H., D.H.B., M.S.D., J.E.C., L.B.T. and R.C. planned the studies. P.G., R.G.B., J.H.E., Q. T., R.N., C.S., T.J., L.A.D., A.K., J.A., J.L., M.E.F., P.A., E.L., S.E., B.G., J.F.S., V.R., T.B., T.C.L., C.H.L., and J.N. conducted experiments, P.G., E.D., R.G.B., J.H.E., C.S., T.J.S., M.J.G., N.V.H., M.S.D., J.E.C. L.B.T. and R.C. interpreted the studies. P.G., J.E.C., and R.C. wrote the first draft of the paper. B.J.D., G.A., S.G.R., D.H.B., M.S.D., and J.E.C. obtained funding. All authors reviewed, edited and approved the paper.

## Data availability

All relevant data are included with the manuscript.

## Supplementary Figures

**Supplementary Figure 1.**
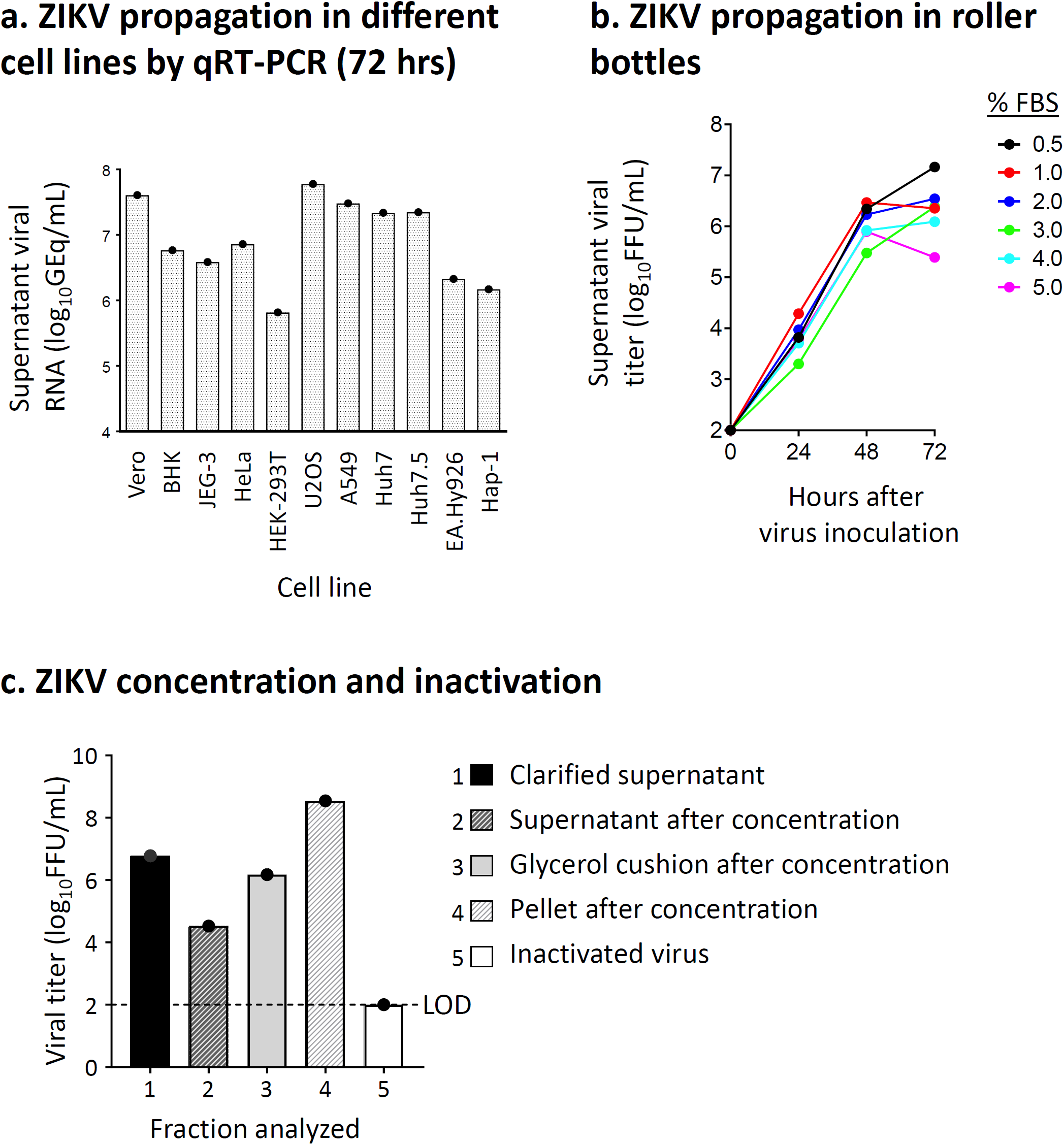
ZIKV stock production. **(a)** Propagation of ZIKV Dakar MA was assessed in a panel of immortalized cell lines. Viral growth was assessed by qRT-PCR from clarified cell culture supernatants harvested 72 hrs after virus inoculation. Data represented a single measurement from one experiment. **(b)** Propagation of ZIKV Dakar MA in Vero cells grown in roller bottles was determined at various FBS concentrations. Viral growth was measured by FFA of culture supernatants. Data represented mean ± SD values of technical duplicates from one experiment. **(b)** Concentration and inactivation of ZIKV Dakar MA was assessed in indicated fractions obtained after ultra-centrifugation of infected Vero cell culture supernatant (72 hrs after virus inoculation), as well as following treatment with hydrogen peroxide. Viral titer was measured by FFA. LOD was 100 FFU/mL. Data represented a single measurement from one experiment.

**Supplementary Figure 2.**
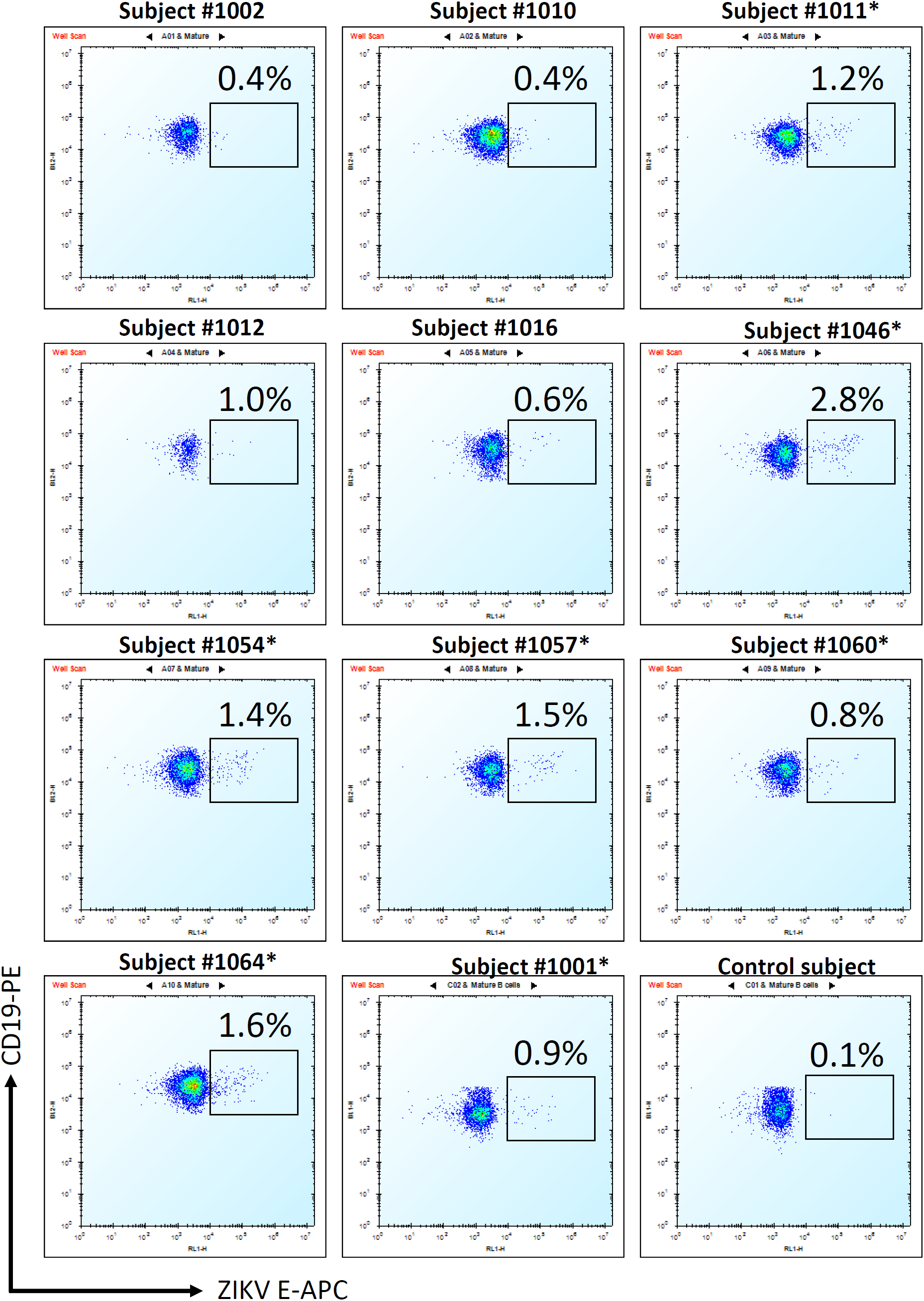
Frequency of ZIKV E-reactive memory B cells identified for previously infected subjects. Percent (%) of ZIKV E-reactive B cells of total B cells was determined by flow cytometric analysis after staining of magnetically enriched B cells from each subject with phenotyping antibodies, biotinylated E protein, and fluorochrome-conjugated streptavidin as detailed in the *Methods* section. Subject with no history for ZIKV exposure served as a control for background E protein staining. Plots show measurements from individual subjects and represent viable CD19^+^IgM^-^IgD^-^ population of magnetically enriched B cells. Seven subjects selected for ZIKV-specific mAb discovery are indicated with “*” symbol.

**Supplementary Figure 3.**
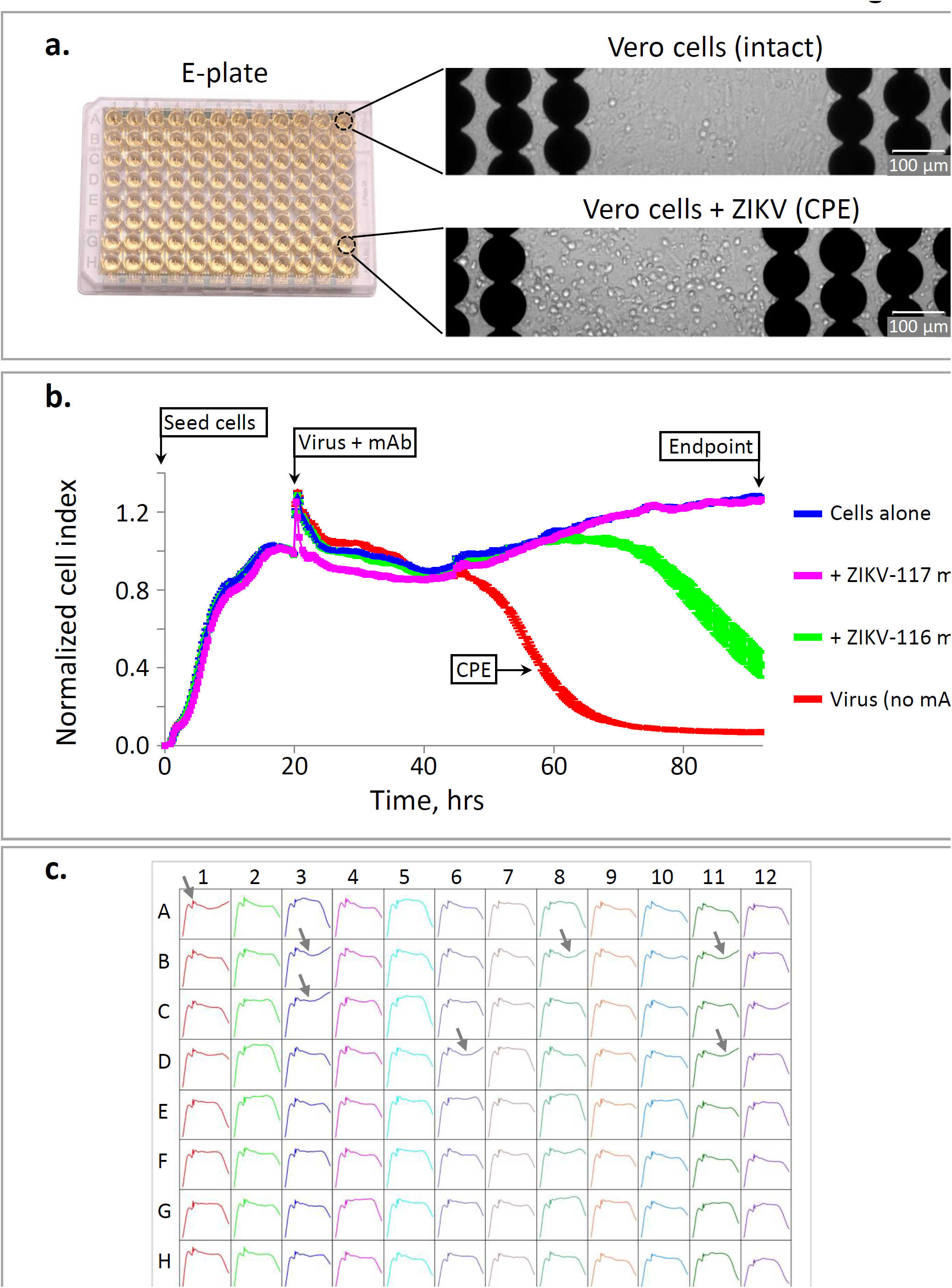
Rapid RTCA screening for neutralizing activity of mAbs to identify lead candidates for *in vivo* protection studies. **(a)** Enlarged brightfield image (left) of shadowed electrodes and adherent Vero cells (with or without virus to visualize CPE) from single wells of 96-well E-plates (right). **(b)** Representative sensorgrams for Vero cells that were inoculated with ZIKV (Brazil strain) in the presence of fully neutralizing ZIKV-117 (magenta) and partially neutralizing ZIKV-116 (green). mAbs ZIKV-117 and ZIKV-116 were described previously ^5^. Uninfected cells (blue) and infected cells without antibody addition (red) served as controls for intact monolayer and full CPE, respectively. Data represented mean ± SD values of technical duplicates. **(c)** Example sensorgrams from one 96-well E-plate analysis showing rapid identification of mAbs that fully neutralize ZIKV (indicated with grey arrows). Neutralization was assessed for 1:25 micro-scale purified mAbs dilution (see *Methods*) using ZIKV Brazil strain. Plates were measured continuously for 45 hrs after applying virus and mAb mixtures to the Vero cell monolayers.

**Supplementary Figure 4.**
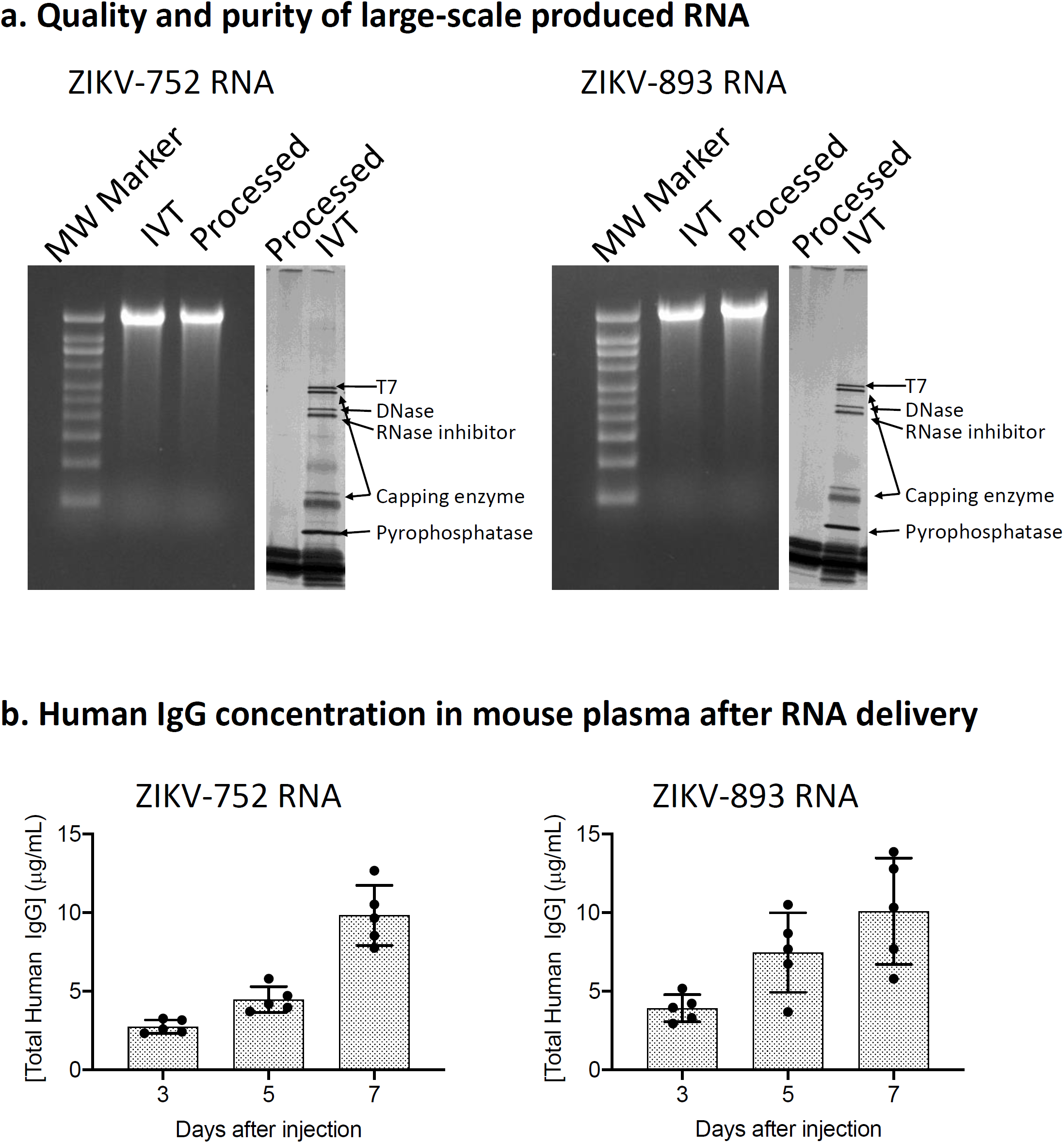
Large-scale production and qualification of ZIKV-753 and ZIKV-893 RNAs. (a) Following large-scale *in vitro* transcription (IVT) and capping, ZIKV-752 and ZIKV-893 RNA were purified and concentrated by Capto Core chromatography and tangential flow filtration, prior to sterile-filtration and storage at −80°C. ZIKV-752 and −893 RNA were analyzed by denaturing gel electrophoresis as well as silver-stain SDS-PAGE to assess quality and purity of post-processed RNA. To verify function, processed ZIKV-752 and −893 were formulated with NLC, and 40 μg of formulation was injected intramuscularly into C57BL/6 mice (n=5/group). Blood was collected on days 3, 5, and 7, and concentrations of total human IgG protein were determined in serum by ELISA using recombinant ZIKV-752 and −893 mAbs to generate a standard curve. Dots show measurements from individual mice and mean ± SD values are shown.

**Supplementary Figure 5.**
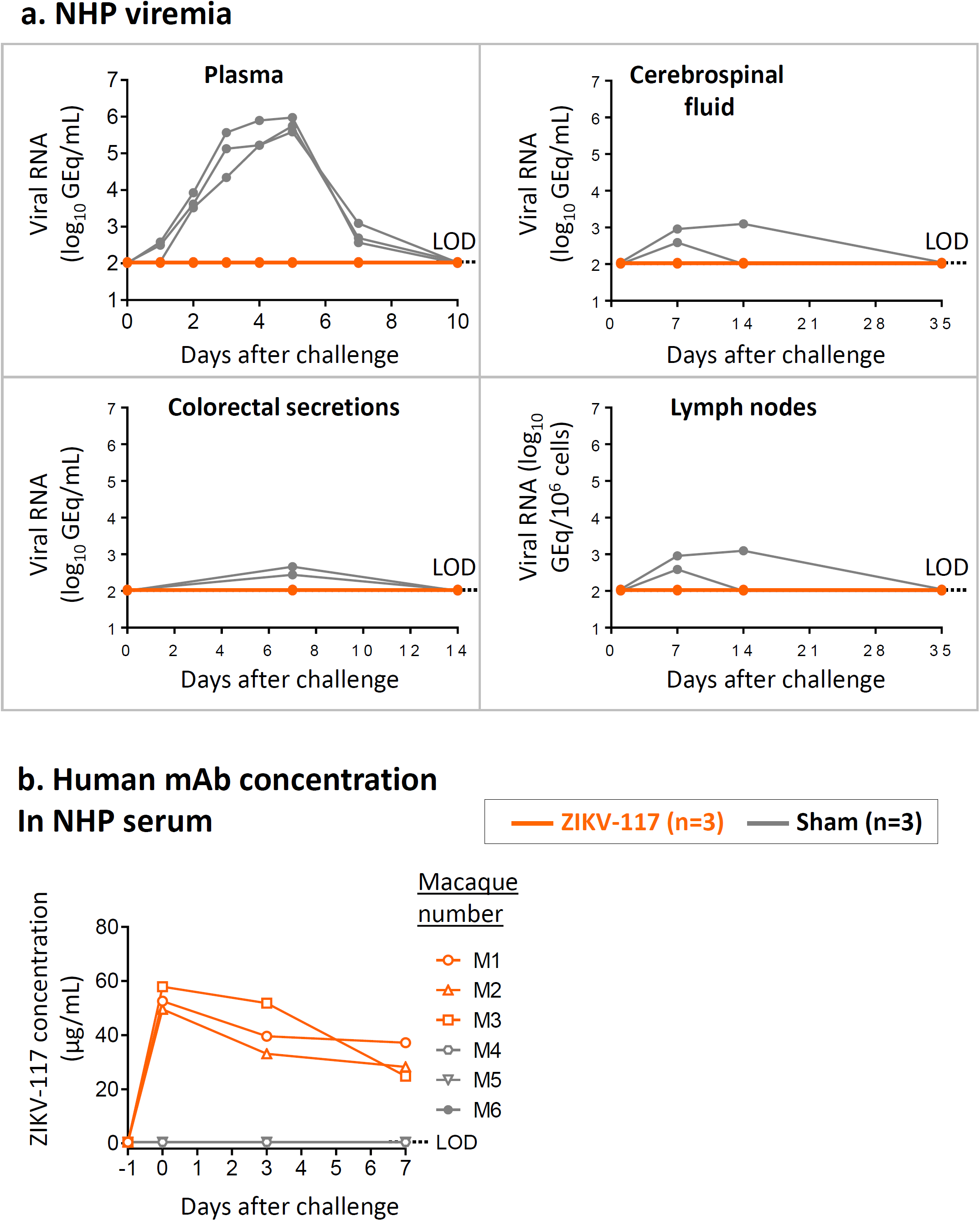
Efficacy of mAb treatment against ZIKV infection in NHPs. Animals received one 10 mg/kg dose of mAb ZIKV-117 (n=3 NHPs) or mAb PGT121 (sham; n=3 NHPs) served as a contemporaneous control intravenously on day −1 and then challenged subcutaneously with a target dose of 10^3^ PFU of ZIKV strain Brazil the next day. (a) qRT-PCR measurement of viremia in plasma and other compartments at indicated time points after virus challenge. (b) Concentration of ZIKV-117 human mAb that was determined in serum of treated and control NHPs at indicated time points after virus challenge.

**Supplementary Table 1.** Research subject demographics and ZIKV exposure history.

| **Subject ID** | **Gender** | **Age** | **Ethnicity** | **ZIKV strain lineage\*** | **Year infected** | **Country in which infection occurred** | **PBMCs collection timepoint** |
| --- | --- | --- | --- | --- | --- | --- | --- |
| 1001 | Male | 41 | Hispanic or Latino | Asian | 2015 | Brazil | 7 months after infection |
| 1011 | Female | 25 | Not Hispanic or Latino | Asian | 2016 | Haiti | March 2016 |
| 1046 | Female | 58 | Hispanic or Latino | Asian | 2016 | Dominican Republic | >90 days after infection |
| 1054 | Female | 62 | Not Hispanic or Latino | Asian | 2016 | Puerto Rico | >90 days after infection |
| 1057 | Male | 40 | Not Hispanic or Latino | Asian | 2016 | Nicaragua | 5 weeks and six months post 1st symptom |
| 1060 | Female | 35 | Not Hispanic or Latino | Asian | 2016 | Guatemala | >90 days after infection |
| 1064 | Male | 19 | Not Hispanic or Latino | Asian | 2016 | Guatemala | >90 days after infection |

\* - The subjects likely had been infected with a strain of Asian lineage of ZIKV, since they contracted the disease during the recent outbreak in South America

**Supplementary Table 3.**
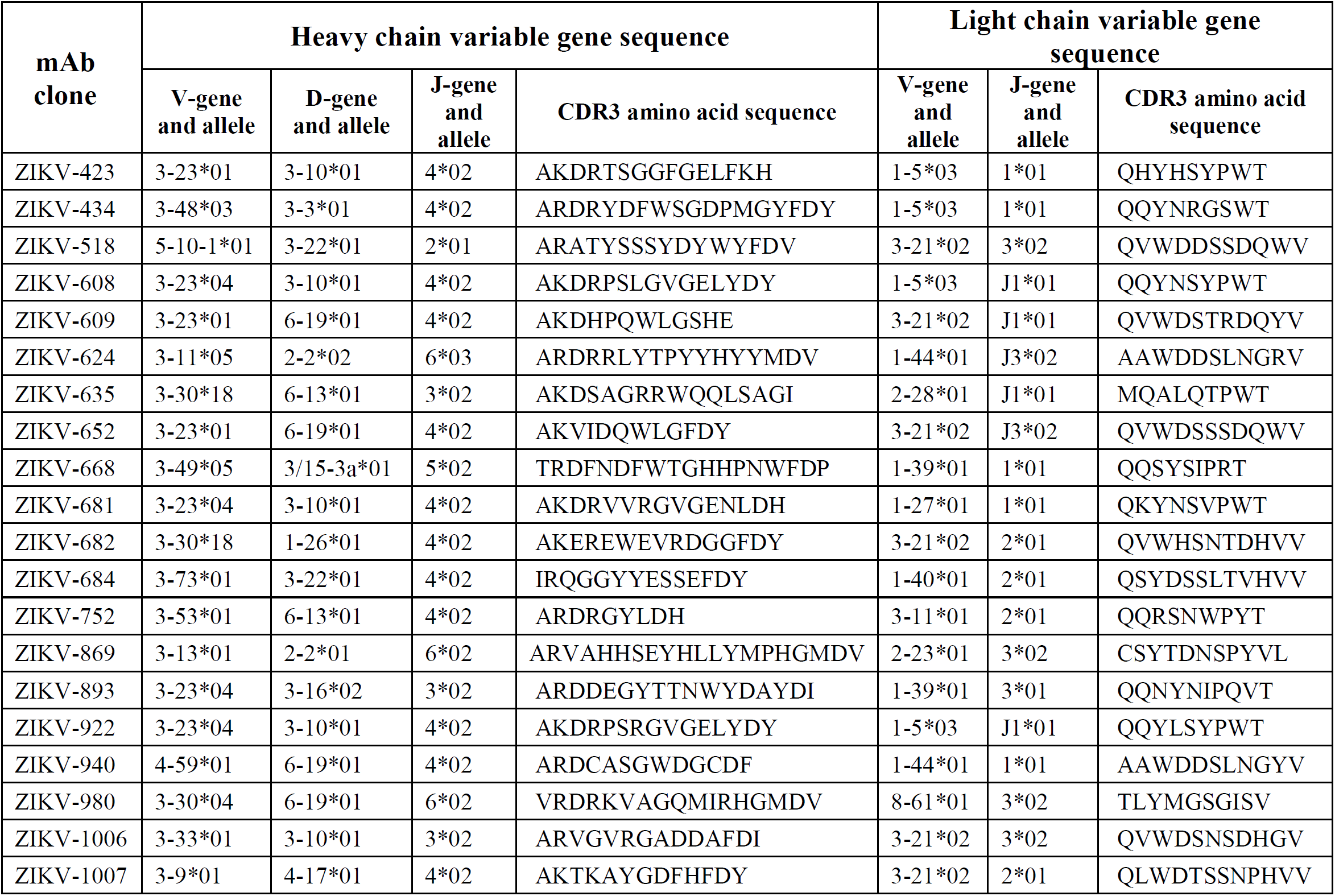
Inferred antibody germline genes and variable region analysis of selected lead candidates that tested *in vivo* in mice

**Supplementary Table 4.**
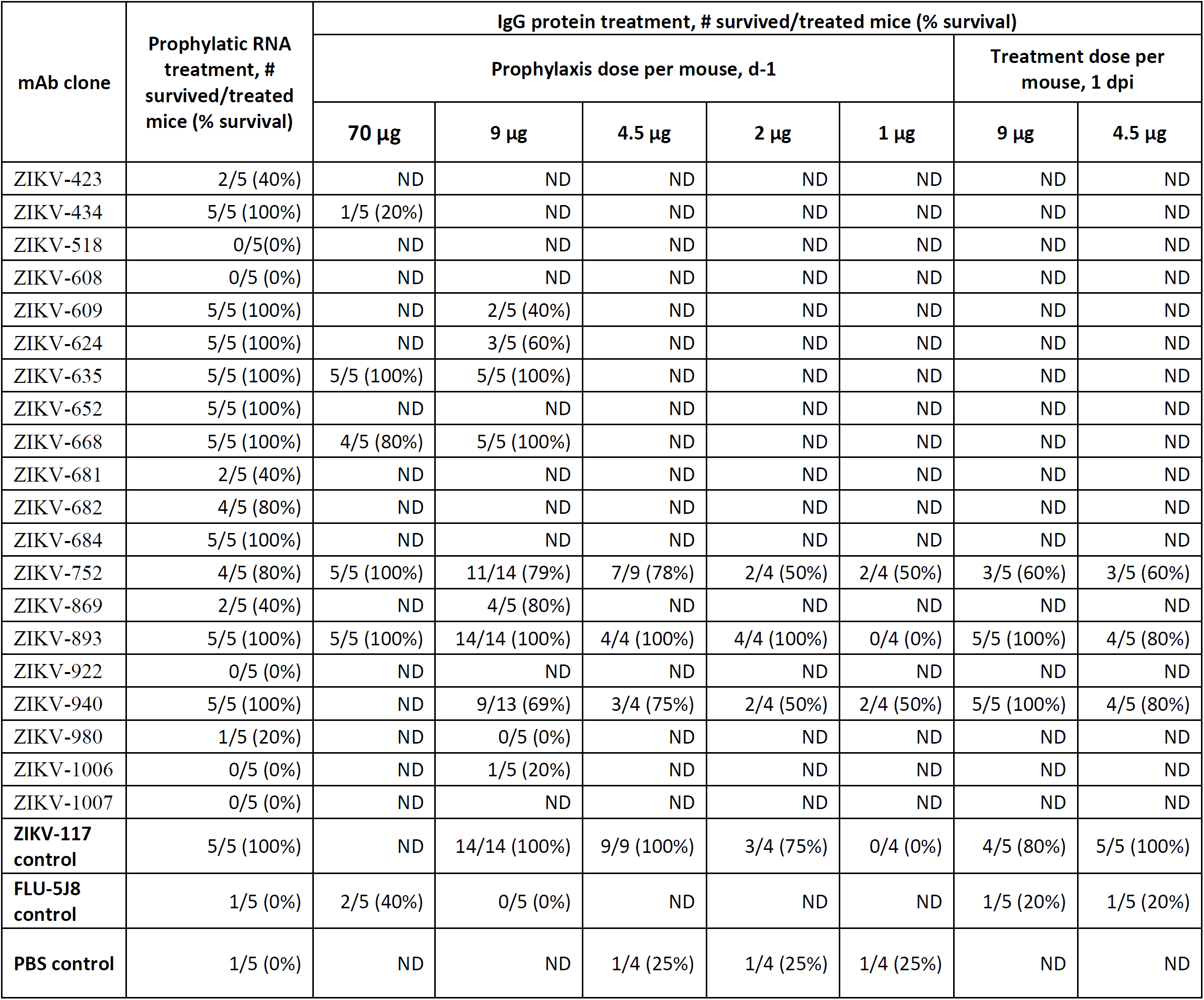
Efficacy of mAb-encoding RNA formulation and IgG protein treatment against ZIKV *in vivo* in mice.

**Supplementary Table 2.**
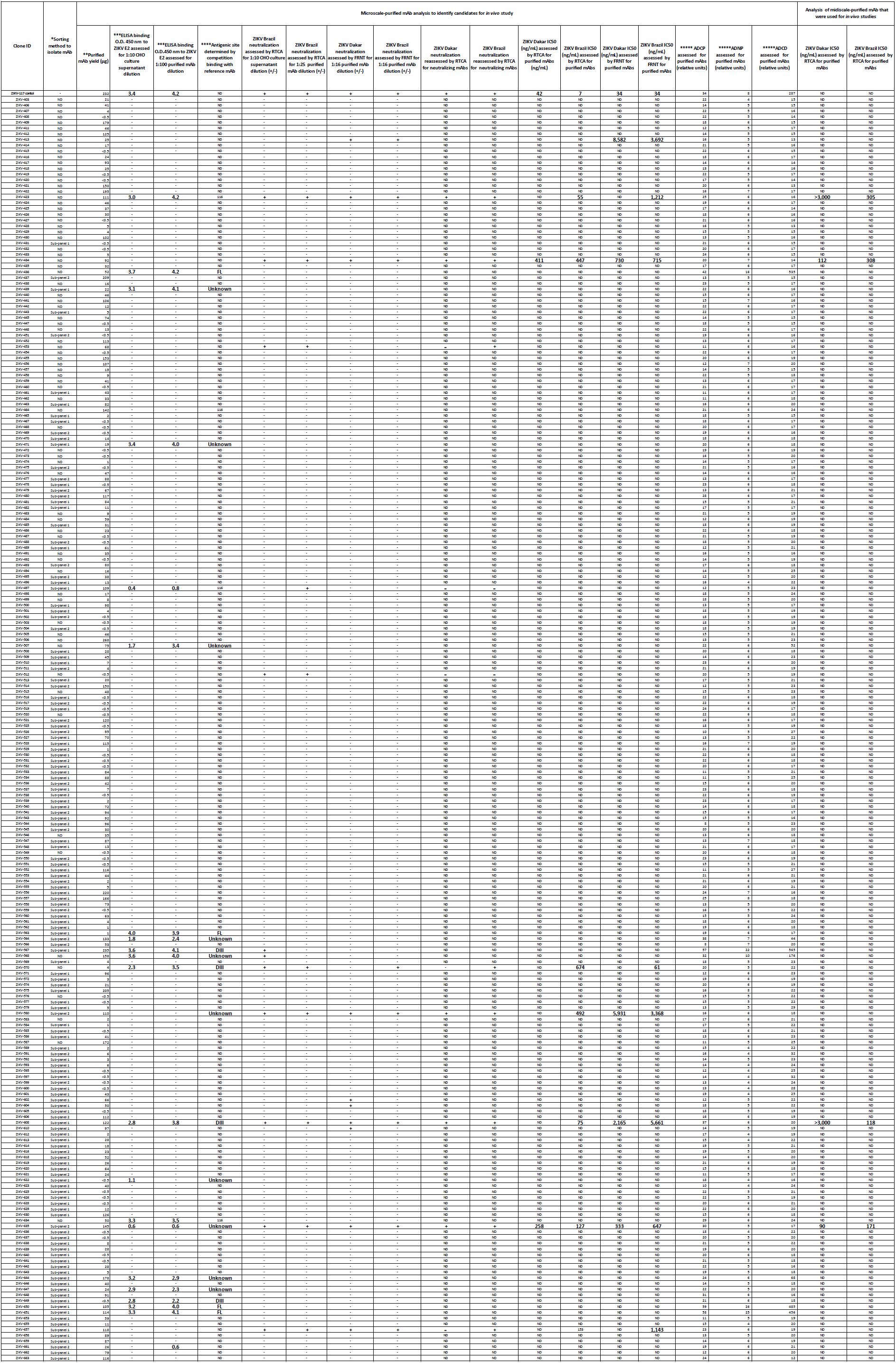
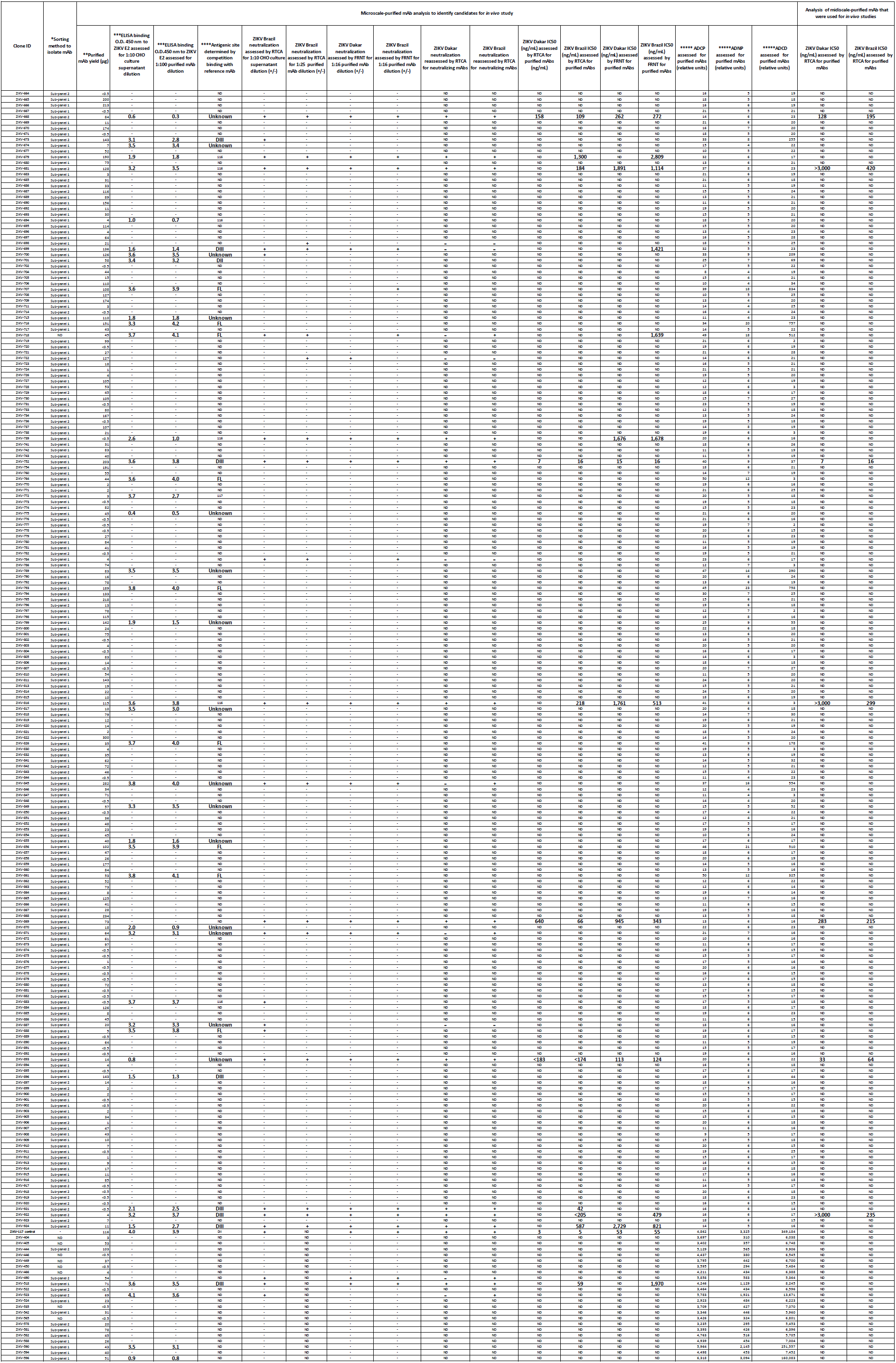
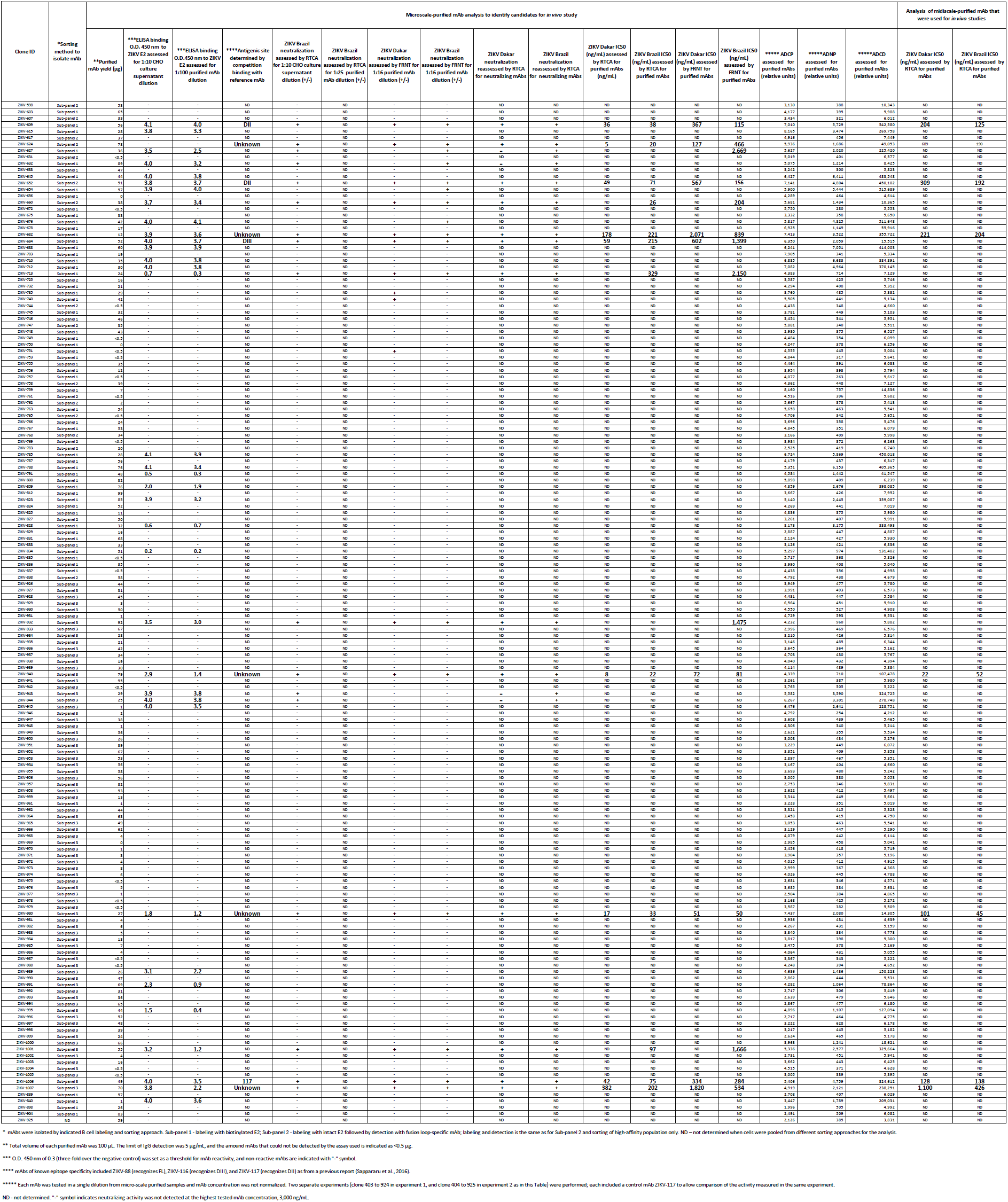
Characteristics of individual mAbs of the 598 recombinant mAbs panel.

